# Profiling dynamic RNA-protein interactions using small molecule-induced RNA editing

**DOI:** 10.1101/2022.06.30.498348

**Authors:** Kyung W. Seo, Ralph E. Kleiner

**Affiliations:** Department of Chemistry, Princeton University, Princeton, NJ, USA 08544

## Abstract

RNA binding proteins (RBPs) play an important role in biology and characterizing dynamic RNA-protein interactions in their native context is essential for understanding RBP function. Here, we develop targets of RNA-binding proteins identified by editing induced through dimerization (TRIBE-ID), a facile strategy for identifying and quantifying state-specific RNA-protein interactions based upon rapamycin-mediated chemically induced dimerization and RNA editing. We perform TRIBE-ID with G3BP1, an abundant RBP and core component of stress granules, to study transcriptome-wide G3BP1-RNA interactions during normal conditions and upon oxidative stress-induced liquid-liquid phase separation (LLPS). We quantify editing kinetics in order to infer interaction persistence and show that stress granule formation strengthens preexisting G3BP1-RNA interactions and induces new RNA-protein binding events. Further, we demonstrate that G3BP1 stabilizes its RNA clients in a dose-dependent manner, suggesting that stress granules function as RNA storage depots. Finally, we apply our method to characterize small molecule modulators of G3BP1-RNA binding. Taken together, our work provides a general approach to profile RNA-protein binding events with temporal control and illuminates the role of LLPS in organizing G3BP1-RNA interactions in the cell.

## INTRODUCTION

RNA-binding proteins (RBPs) play an important role in the post-transcriptional regulation of gene expression^1^. Over 1000 proteins have been annotated as RBPs and they can affect diverse processes throughout the RNA life cycle including splicing, nuclear export, localization, translation, and metabolism^1^. Since RBPs typically bind and regulate multiple RNA transcripts, transcriptome-wide identification and characterization of cellular RBP-RNA interactions, or ribonucleoproteins (RNPs), provides important insights into the biological function of RBPs. Further, dysregulation of RBP-RNA interactions has been linked to human disease and modulating these interactions with small molecules and oligonucleotides has emerged as a promising therapeutic strategy^2,3^. Thus, profiling the native RNA targets of individual RBPs is important for understanding molecular mechanisms of gene expression regulation and disease phenotypes^4^.

Many RNP complexes exist as higher-order assemblies within cells. Recent evidence has indicated that the formation of RNA-protein granules or condensates is driven by liquidliquid phase separation (LLPS), and condensate formation can be a dynamic and highly regulated process^5^. The physical forces governing RNA-protein condensate formation have been studied *in vitro* and in cells and involve the sum of homotypic and heterotypic RNA and protein interactions that can be modulated by protein and RNA structure and sequence, macromolecule concentration, and post-translational and post-transcriptional modifications, among other factors^5^. While studies of RNA-protein condensate formation have provided a biophysical framework for understanding this process, the fate of individual protein-RNA interactions during biomolecular phase separation and the effect of condensate formation on the biological function of RNP complexes remains poorly understood. This is due in large part to the challenge of characterizing dynamic RNP interactions within the cell.

Ras GTPase-activating protein (SH3 domain)-binding protein 1 (G3BP1) is an abundant RBP that can regulate RNA metabolism and translation^6^. G3BP1 has diverse biological functions and has been linked to tumor development, immune activation, stress response, and neuronal development and activity^6^. Much of the recent work on G3BP1 has focused on its role in the formation of stress granules, cytoplasmic RNA-protein condensates that assemble via LLPS in response to a variety of stresses^7^. Prevailing models for G3BP1-mediated stress granule formation involve protein dimerization and sequence-independent RNA binding, with G3BP1 proposed to function as the central node in a diverse network of RNA and protein interactions^8–11^. Despite these advances in our understanding of G3BP1-mediated stress granule-assembly, there remains a major gap in the characterization of individual G3BP1-RNA interactions in non-stressed cells and their persistence during stress granule formation, as well as the effect of G3BP1 binding on RNA metabolism and translation. G3BP1-RNA interactions have been profiled in non-stressed cells^12–14^, but independent datasets have failed to reach consensus, and G3BP1-RNA binding has been proposed to be both RNA sequence-dependent^12,15^ and independent^10,13^ In addition, we lack information on G3BP1-RNA interactions within stress granule condensates, and conflicting data exist regarding the role of G3BP1 in RNA metabolism with reports of substrate stabilization^12,16^ and destabilization^17^.

The most common strategies for identifying RNA targets of an RBP rely upon photoinduced crosslinking and immunoprecipitation coupled with high-throughput RNA sequencing (CLIP-seq or HITS-CLIP, and many variations thereof)^18–20^. While CLIP approaches provide a general method to characterize native cellular RNA-protein interactions at nucleotide resolution^18–20^, there are a number of challenges that have limited its usage including high input requirements, antibody availability, lack of UV penetration into tissue samples, and the time and labor-intensive nature of the protocol. Further, CLIP is generally not suitable for characterizing interaction strength or residence time and variability in experimental methods and bioinformatic analysis can have a significant impact on detected transcripts, leading to, in some cases, poor reproducibility among CLIP datasets for the same RBP^18^.

To address the limitations of CLIP, complementary strategies for RBP-RNA interaction analysis are in development. In particular, one class of approaches relies upon fusing RNA modifying enzymes to the RBP of interest in order to ‘mark’ substrate transcripts with post-transcriptional RNA modifications that can be detected by sequencing^21^. Notable examples of this strategy include TRIBE (targets of RNA-binding proteins identified by editing)^22^, RNA tagging^23^, and STAMP (surveying targets by APOBEC-mediated profiling)^24^, which use adenosine deaminase, poly(U) polymerase, and cytidine deaminase, respectively. These strategies are attractive since they do not require extensive biochemical steps or large amounts of cells, however, their generality has not been investigated broadly. Additionally, while CLIP-based methods can analyze dynamic processes, enzymatic labeling methods lack temporal resolution due to asynchronous expression of the RBP-enzyme fusion in a cellular population. Therefore, while enzymatic labeling methods may provide a facile and efficient approach to characterize the RNA substrates of an RBP, they are not suitable for studying dynamic interactions found in biology.

Here, we develop TRIBE-ID (targets of RNA-binding proteins identified by editing induced through dimerization; Fig. 1a), a method for profiling and quantifying dynamic RNA-protein interactions in live cells with temporal precision. TRIBE-ID relies upon ADAR-mediated A-to-I editing to mark RBP substrate transcripts and rapamycin-induced FRB/FKBP dimerization^25,26^ to control the timing of editing. We apply TRIBE-ID to profile cytoplasmic G3BP1-RNA interactions and G3BP1-RNA binding within stress granules, and quantify interaction persistence by measuring the accumulation of A-to-I modifications on G3BP1 substrates over time. We identify RNA transcripts involved in high-persistence interactions that are strongly stabilized by G3BP1 binding in cells and exhibit lower translation efficiency. Further, we show that the hierarchy of G3BP1-RNA interactions in non-stressed cells is conserved during stress granule formation and demonstrate a global increase in G3BP1-RNA binding and G3BP1 substrates during biomolecular phase separation. Finally, we demonstrate that TRIBE-ID can be used to characterize the effects of small-molecule G3BP1 modulators upon global G3BP1-RNA interactions. Taken together, our work provides a robust method to identify and quantify state-specific RBP-RNA interactions and screen small molecule RBP-RNA inhibitors, illuminates dynamic G3BP1-RNA interactions during the integrated stress response, and establishes G3BP1 as a global post-transcriptional regulator.

**Figure 1.**
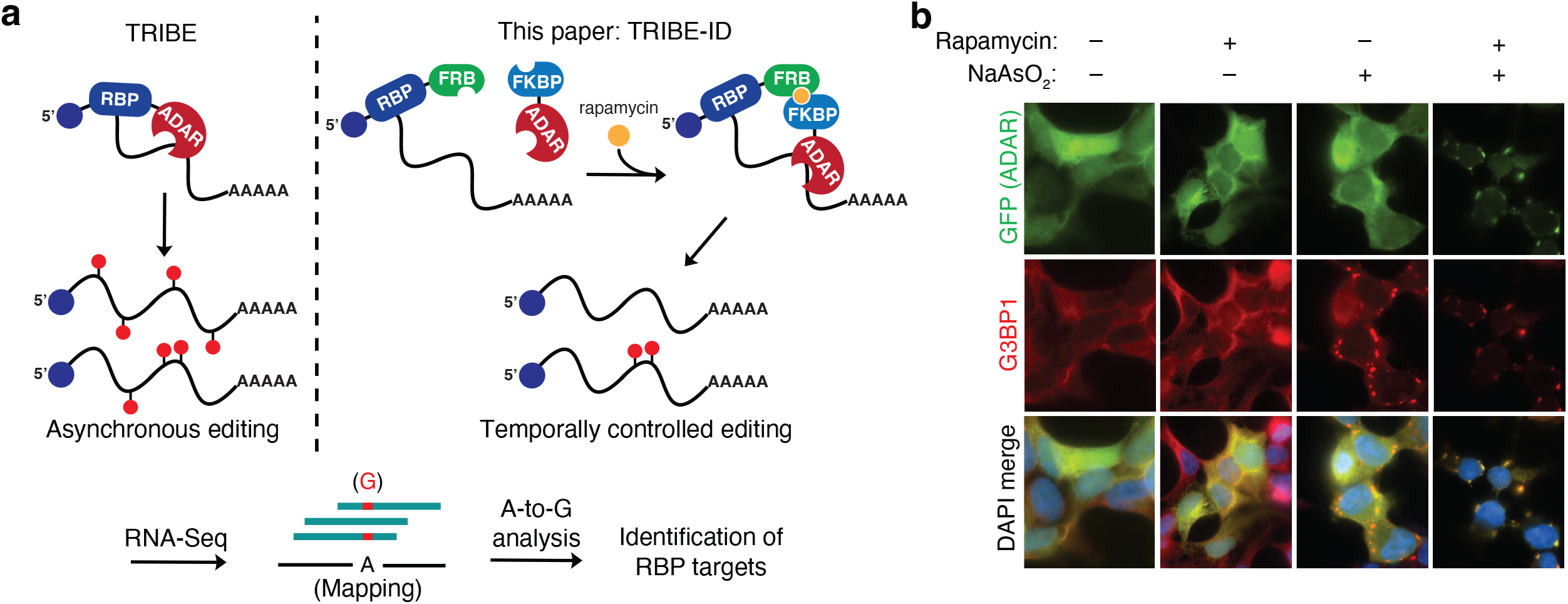
Temporally controlled RNA-protein interaction analysis. **(a)** Workflow for RNA-protein interaction analysis using TRIBE/HyperTRIBE or TRIBE-ID. **(b)** Localization of G3BP1-FRB and FKBP-hADAR(E488Q)-GFP constructs in response to rapamycin treatment and NaAsO_2_ stress. Imaging was performed using immunofluorescence microscopy.

## RESULTS

### Rapamycin-mediated dimerization of G3BP1-FRB and FKBP-ADAR

To develop our method (Fig. 1a), we designed RBP and ADAR protein constructs that would assemble and modify RBP substrate transcripts only in the presence of a small molecule. We chose the well-established FRB-FKBP chemically induced dimerization (CID) system that is responsive to rapamycin and related derivatives^27^. This system has been utilized widely for CID in diverse biological contexts, relies on small protein tags that are unlikely to perturb native function, and has been used successfully for small molecule-dependent nucleic acid editing^27,28^. In contrast to the canonical TRIBE method where expression of an RBP-ADAR fusion protein mediates constitutive editing^22^, we envisioned that the co-expression of separate RBP and ADAR proteins containing complementary FRB/FKBP dimerization tags would enable conditional RNA editing in a rapamycin-dependent manner, allowing temporal control over RBP-RNA interaction analysis. In order to explore this strategy, we chose the stress granule-associated RBP G3BP1. While G3BP1-RNA interactions have been studied using CLIP methods^12–14^, these analyses have not converged upon a consensus G3BP1-RNA interactome. Further, G3BP1 subcellular localization can be rapidly modulated by stress resulting in its accumulation in stress granules^7^, and the effect of stress granule formation on global G3BP1-RNA interactions is not well understood. Therefore, we envisioned that stress-induced G3BP1 relocalization would provide a dynamic system for studying chemically induced protein dimerization and state-specific small-molecule dependent RNA editing.

To investigate rapamycin-induced dimerization between ADAR and G3BP1, we performed immunofluorescence microscopy. First, we generated constructs with FKBP fused to the N-terminus of human ADAR2 catalytic domain (hADAR) and FRB fused to the C-terminus of G3BP1 and co-expressed them in HEK293T cells. Next, we evaluated the subcellular localization of G3BP1-FRB and FKBP-hADAR in the presence or absence of stress. In the absence of stress, G3BP1 and ADAR constructs showed diffused cytosolic localization (Fig. 1b). Gratifyingly, when cells were treated with 100 nM rapamycin, we observed overlap between G3BP1 and FKBP-hADAR constructs as early as 30 minutes post-treatment. This was most apparent when we combined rapamycin with sodium arsenite treatment and visualized clear accumulation of FKBP-ADAR and G3BP1 in stress granules (Fig. 1b). In contrast, sodium arsenite treatment alone results in G3BP1 recruitment to stress granules whereas FKBP-ADAR remains diffused throughout the cytoplasm. Taken together, our immunofluorescence microscopy data demonstrates robust rapamycin-mediated CID between FKBP-ADAR and G3BP1-FRB in cells.

### G3BP1 TRIBE analysis with human and Drosophila ADAR2 catalytic domains

To benchmark the TRIBE-ID method and optimize detection of bona fide G3BP1 substrates, we evaluated different ADAR catalytic domains in isolation and constitutively fused to G3BP1 (i.e. the canonical TRIBE/HyperTRIBE method^22,29^). An ideal ADAR should display high editing activity when dimerized or fused to G3BP1 and low background editing when expressed alone or without dimerization. We chose three ADARs for comparisons: 1) *Drosophila* ADAR catalytic domain with hyperactive E488Q mutation (dADAR(E488Q)) used in the HyperTRIBE method^29^; 2) human ADAR2 catalytic domain with E488Q mutation (hADAR(E488Q)); and 3) human ADAR2 catalytic domain with E488Q and T375G mutations (hADAR(E488Q/T375G), which was previously used for single-site CRISPR/Cas13-mediated RNA editing^30^ (Fig. 2a). We generated fusions of each ADAR and G3BP1, and transfected HEK293T cells with plasmids encoding G3BP1-ADAR or ADAR alone. After ADAR expression, we harvested cells, isolated poly(A)RNA, and performed Illumina sequencing. A-to-I editing sites in each sample were identified using the TRIBE bioinformatic pipeline comparing against untransfected HEK293T cells (Fig. 2b). Two independent biological replicates were used for all conditions.

**Figure 2.**
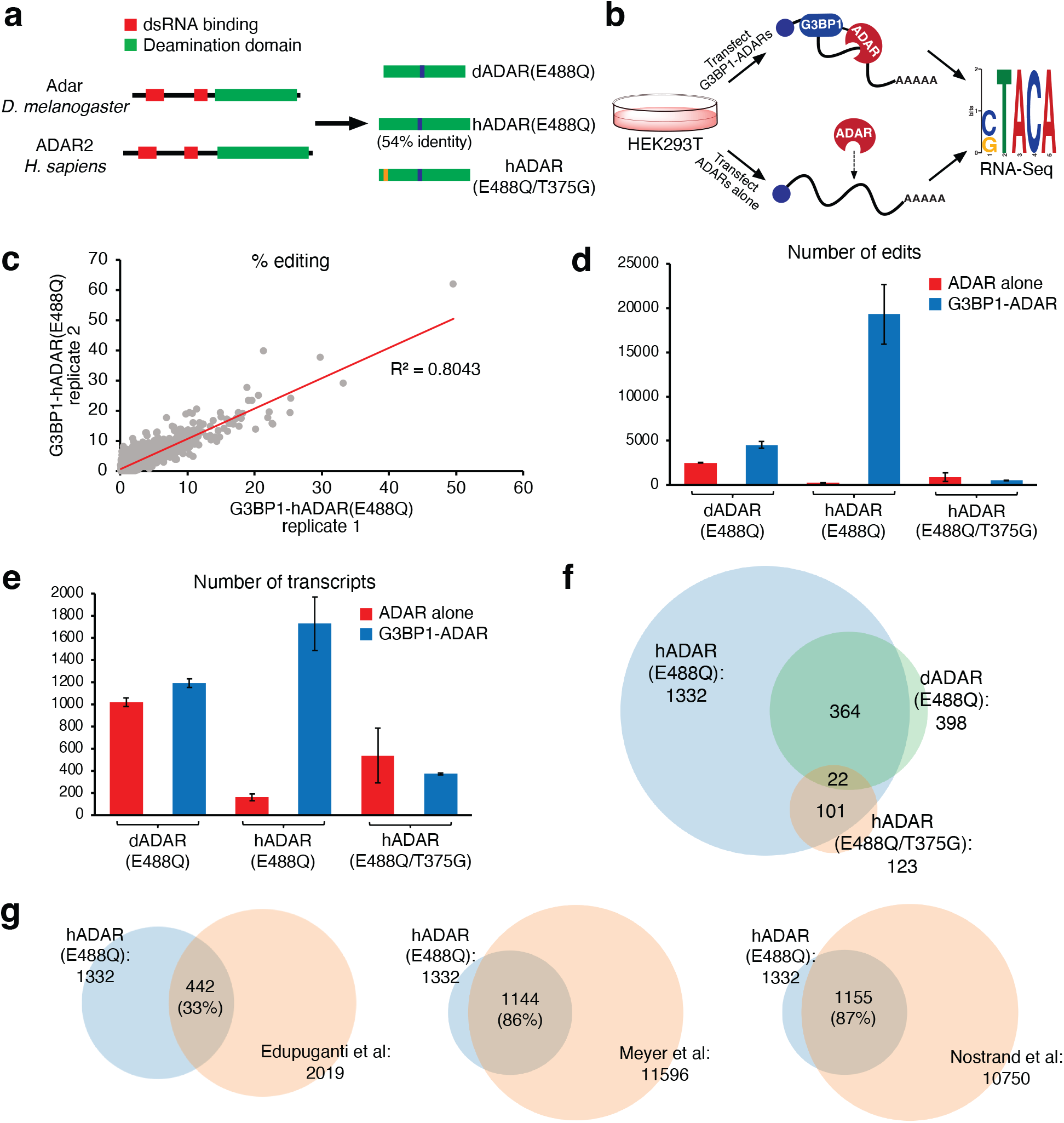
HyperTRIBE analysis of G3BP1-RNA binding. **(a)** ADAR2 catalytic domains used in this work. **(b)** Workflow for screening G3BP1-ADAR and ADAR constructs. **(c)** RNA editing reproducibility for G3BP1-hADAR(E488Q) in two independent biological replicates. **(d)** Average number of edit sites detected in cells transfected with G3BP1-ADAR or ADAR. **(e)** Average number of edited transcripts detected in cells transfected with G3BP1-ADAR or ADAR. **(f)** Venn diagram showing overlap between transcripts detected with each G3BP1-ADAR construct. **(g)** Venn diagrams showing overlap between G3BP1-hADAR(E488Q) targets and previous CLIP data.

Comparative editing site analysis allowed us to differentiate G3BP1-dependent editing from background ADAR editing. RNA editing events in G3BP1-ADAR transfected samples were reproducible in their location and frequency (Fig. 2c and Supplementary Fig. 1; R^2^ = 0.62-0.81). The reproducibility was considerably lower (R^2^ = 0.13-0.45) in samples transfected with ADAR alone, likely due to the non-specific nature of these interactions (Supplementary Fig. 1). In G3BP1-hADAR(E488Q) expressing cells, we identified an average of nearly 20,000 edit sites across replicates (Fig. 2d), whereas we detected only 238 edit sites in cells containing hADAR(E488Q) alone, indicating that the overwhelming majority of editing is mediated by G3BP1-dependent RNA binding. In cells transfected with G3BP1-dADAR(E488Q) or G3BP1-hADAR(E488Q/T375G), we detected ~4,500 and ~900 edit sites, respectively, considerably fewer than with G3BP1-hADAR(E488Q), and background editing from expression of dADAR(E488Q) or hADAR(E488Q/T375G) alone was higher than with hADAR(E488Q) (Fig. 2d). The number of edited transcripts (as opposed to individual edit sites) showed similar trends across all three constructs, with G3BP1-hADAR(E488Q) editing the highest number of transcripts (~1700) and hADAR(E488Q) alone showing the lowest background transcript editing (Fig. 2e).

In order to annotate G3BP1 RNA substrates identified using each construct, we compared G3BP1-ADAR edit sites and those edited by the respective ADAR alone, taking only those unique to G3BP1-ADAR expression as bona fide G3BP1-associated RNA transcripts. We also eliminated transcripts with only intronic edit sites^31^ (since G3BP1 is primarily localized to the cytoplasm) and restricted our analysis to edited transcripts found in both biological replicates^22^. Using this analysis, we identified 1332 G3BP1 substrates with G3BP1-hADAR(E488Q), but only 398 and 123 substrates using G3BP1-dADAR(488Q) and G3BP1-hADAR(E488Q/T375G), respectively. Further, the majority of G3BP1 substrates identified using dADAR(488Q) or hADAR(488Q/T375G) were also found in the hADAR(E488Q) dataset (Fig. 2f), suggesting that all enzymes have overlapping substrate scope and that hADAR(E488Q) has the highest sensitivity.

Since TRIBE analysis of human G3BP1 substrates has not been previously reported, we compared the 1332 transcripts identified with G3BP1-hADAR(E488Q) against reported G3BP1-CLIP targets^12–14^ (Fig. 2g). Three CLIP studies have reported G3BP1 transcripts: Edupuganti *et al*.^12^ and Meyer *et al.^13^* performed PAR-CLIP, and Van Nostrand *et al*.^14^ performed eCLIP. We found a high degree of overlap (86-87%) between our 1332 G3BP1 TRIBE targets and the CLIP datasets from Meyer *et al*. or Van Nostrand *et al*., respectively (Fig. 2g). In contrast, only 33% of G3BP1-hADAR(E488Q) targets were found in the CLIP data from Edupuganti *et al*.^12^; it is important to note that ~5-fold fewer peaks and transcripts were detected in this dataset compared to the other two G3BP1 CLIP datasets. We also performed RIP-Seq using FLAG-tagged G3BP1 and found that 834 (62.6%) of G3BP1-hADAR(E488Q) targets were detected (IP/input > 1, P-adj < 0.05) (Supplementary Fig. 2). Analysis of sequences surrounding edit sites in G3BP1-hADAR(E488Q) TRIBE data showed enrichment of a number of motifs (Supplementary Fig. 3). Two motifs were also found in cells expressing hADAR(E488Q) alone and are likely to represent preferred ADAR2 substrate sequences (Supplementary Fig. 3). Interestingly, we also found a previously described G3BP1-associated CUGGA motif^13^ as one of the top enriched motifs in our data (Supplementary Fig. 3). Gene ontology (GO)-term enrichment analysis showed significant enrichment in mRNA metabolism, translation, and cell cycle, which were also significantly enriched in previous G3BP1 studies^12–14^ (Supplementary Fig. 4). Taken together, our results indicate that TRIBE can efficiently identify G3BP1 substrates in human cells, and demonstrate that hADAR(E488Q) has the lowest background editing and highest sensitivity of the ADAR constructs that we evaluated. Therefore, we utilized hADAR(E488Q) for all subsequent TRIBE-ID experiments.

### Temporally controlled G3BP1-RNA interaction analysis with TRIBE-ID

Next, we used rapamycin-dimerizable FKBP-hADAR(E488Q) and G3BP1-FRB for temporal editing of G3BP1 targets (Fig. 3a). While reported TRIBE and HyperTRIBE editing experiments typically express the RBP-ADAR fusion protein in cells for 24 hours, ADAR editing of RBP targets can occur more rapidly^32^, and we expected that short treatments with rapamycin would likely result in detectable substrate editing. Therefore, we treated cells that stably express G3BP1-FRB and FKBP-hADAR(E488Q) with rapamycin and harvested mRNA for analysis after 2, 4, or 8 hours (Fig. 3a). The sequencing depth across all samples was kept comparable (Supplementary Table 1). As a control for background A-to-I editing, we used the same cell line expressing FKPB/FRB constructs but without rapamycin treatment, which showed similarly low levels of background editing (~100 background editing events) as observed in cells transiently expressing hADAR(E488Q) alone (Supplementary Fig. 5). In contrast, after 2 hr of rapamycin treatment, we detected ~2000 editing events distributed over ~700 transcripts (Fig. 3b and 3c). The number of edits increased with longer rapamycin treatment reaching ~3800 editing events on ~1300 transcripts after 8 hr treatment. Similar to our analysis of the G3BP1 TRIBE experiment, we restricted our analysis of TRIBE-ID data to edited transcripts found in both biological replicates, and identified 483, 547, and 903 transcripts at 2, 4, and 8 hr time points, respectively (Fig. 3d). Among these, 287 transcripts were detected at all three time points, suggesting that these transcripts are frequent, high-confidence substrates of G3BP1. While we detected fewer G3BP1 substrates using TRIBE-ID than with TRIBE, likely due to shorter editing time and incomplete rapamycin-mediated dimerization, there was substantial overlap between transcripts identified using both approaches with 60-66% of transcripts found at each time point using TRIBE-ID also identified in the G3BP1 TRIBE experiment, and similar enrichment of GO terms (Fig. 3d and Supplementary Fig. 6). Moreover, we performed motif analysis on sequences surrounding the editing sites of 287 transcripts found at all three time points (Fig. 3e) and identified similar motifs shared between TRIBE-ID and TRIBE data for G3BP1 (Supplementary Fig. 3). Taken together, our data indicates that TRIBE-ID can identify native G3BP1-RNA interactions with resolution as low as 2 hr.

**Figure 3.**
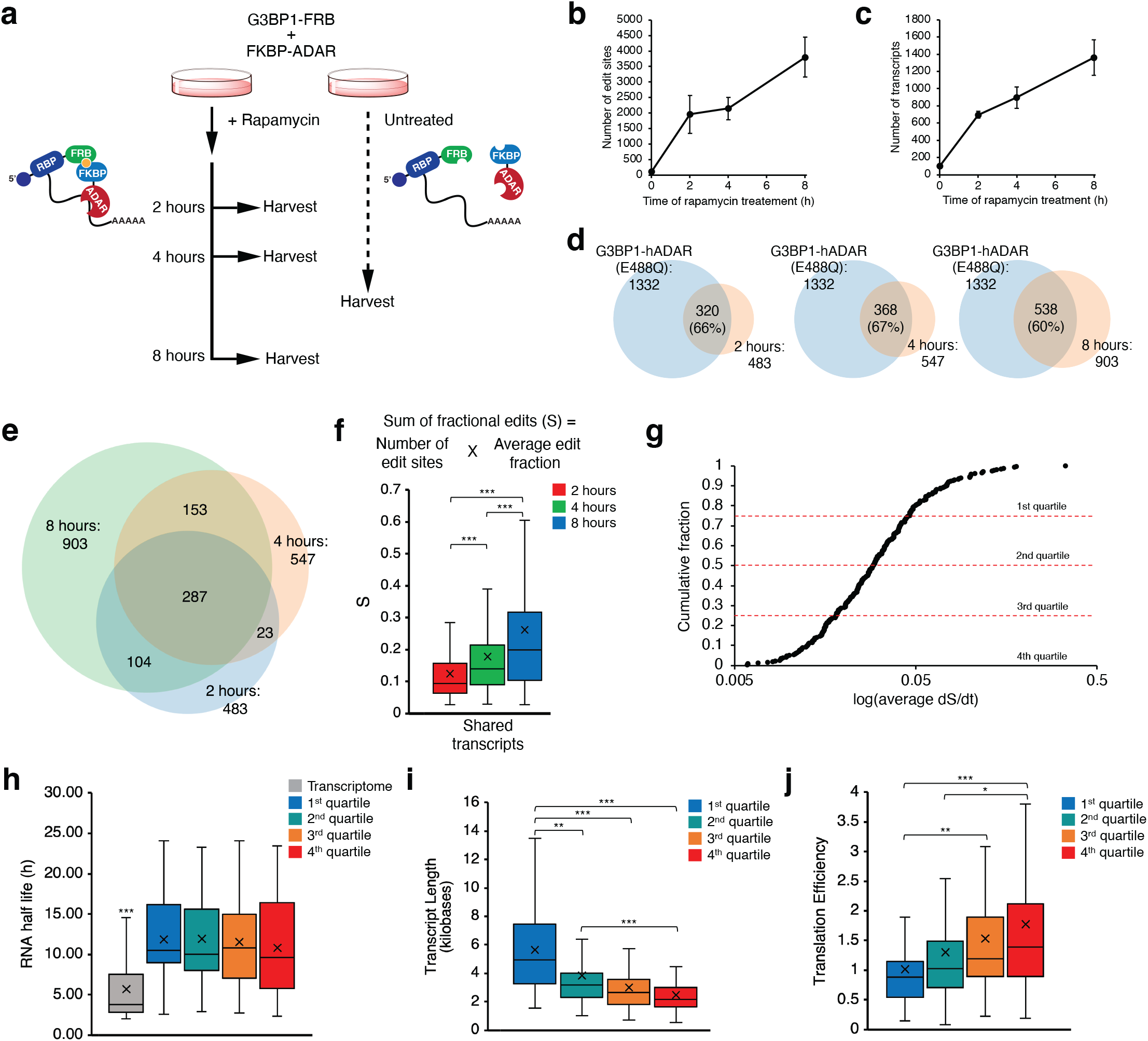
TRIBE-ID detects G3BP1-RNA interactions with temporal control. **(a)** Workflow for G3BP1 TRIBE-ID. **(b)** Average number of edit sites detected with TRIBE-ID across time points. **(c)** Average number of edited transcripts detected with TRIBE-ID across time points. **(d)** Venn diagrams showing overlap between transcripts detected by G3BP1-hADAR(E488Q) (TRIBE) and those detected by TRIBE-ID at each time point of rapamycin treatment. **(e)** Venn diagram showing overlap between transcripts detected using 2, 4, or 8 hours of rapamycin treatment. **(f)** Sum of fractional edits (S) calculation and boxplot depicting S of 287 transcripts shared across all three G3BP1 TRIBE-ID time points. “x” indicates the mean, and the middle line indicates the median. **(g)** Cumulative distribution of G3BBP1 TRIBE-ID transcripts identified at all three time points ranked by their dS/dt values. Dotted red lines indicate quartiles. **(h)-(j)** Boxplots depicting the half-life **(h)**, transcript length **(i)**, and translation efficiency **(j)** of transcripts in different dS/dt quartiles. “x” indicates the mean, and the middle line indicates the median. *: p < 0.05; **: p < 0.005; ***: p < 0.0005

### Quantifying G3BP1-RNA association with TRIBE-ID

In addition to an increase in the number of identified RNA substrates with longer rapamycin treatment, we also expected that transcripts identified at multiple time points would accumulate A:I edit events as a function of their association time with G3BP1, assuming that editing sites are not limiting. This could manifest as an increase in the number of editing sites across a substrate transcript or an increase in editing stoichiometry. To investigate these processes, we analyzed the 287 G3BP1 substrate transcripts found at 2 hr, 4 hr, and 8 hr rapamycin treatment (Fig. 3e). We defined a parameter, S, as the sum of all editing fractions (Fig. 3f), which corresponds to the average editing stoichiometry found across all editing sites multiplied by the number of editing sites, and computed S for each transcript at 2 hr, 4 hr, and 8 hr. Gratifyingly, we observed a statistically significant increase in the average S value of the 287 shared transcripts over time, indicating that editing events accumulate on RBP substrate transcripts in a time-dependent fashion (Fig. 3f). Moreover, 153 of 287 transcripts showed an increase in editing across all time points. Since the number of available editing sites on an RNA could depend upon its sequence composition and length, we chose to focus on the change in S over time (dS/dt) rather than the magnitude of S, which we hypothesized would reflect the residence time of G3BP1 on its RNA substrates. Transcripts were ranked according to average dS/dt and grouped into quartiles for further analyses (Fig. 3g). We compared the transcripts in the top quartile against reported G3BP1-CLIP data^12–14^ and found 59%, 86%, and 97% were detected in Edupuganti *et al*.^12^, Meyer *et al.^13^*, and Van Nostrand *et al*^14^ data, respectively (Supplementary Fig. 7). This degree of overlap is similar to or higher than observed with the 287 transcripts shared across all time points in TRIBE-ID or the targets identified using TRIBE (Fig. 2g, Supplementary Fig. 7), suggesting that transcripts with higher dS/dt values correlate with higher confidence or higher persistence G3BP1-RNA interactions.

### Properties of G3BP1 substrate RNAs

G3BP1 has been demonstrated to regulate the turnover of target mRNAs^8^. While some studies have shown that G3BP1 promotes stabilization of its targets^12,16^, others have shown that G3BP1 can accelerate RNA turnover^17^. Therefore, we compared our TRIBE-ID data against previously reported mRNA half-life data^33^. We found that transcripts shared across all three TRIBE-ID timepoints have roughly 2-fold longer half-life than the average transcriptome-wide value (Fig. 3h). This observation was most apparent for transcripts in the top dS/dt quartile, which exhibited 31% lower variance from the mean than those in the bottom quartile (Fig. 3h). In addition, we found that transcript length positively correlated with dS/dt (Fig. 3i). While longer transcripts are likely to have more available editing sites, we did not observe a significant correlation between number of edit sites and transcript length at any of the time points investigated in our G3BP1 TRIBE-ID analysis (Supplementary Fig. 8) or in our analysis of background editing in cells expressing hADAR(E488Q) alone (Supplementary Fig. 9). Therefore, we favor a model where the correlation of dS/dt with transcript length is likely driven by the presence of additional G3BP1 binding sites on these RNAs. Finally, we studied the translation efficiency of G3BP1 substrates using available genome-wide ribosome profiling data^34^. We found dS/dt negatively correlated with translational efficiency (Fig. 3j). G3BP1 has been shown to negatively regulate translation in neurons^35^, and our findings indicate that G3BP1 association is negatively correlated with translation efficiency. Together, G3BP1 binding appears to be positively correlated with stability and transcript length, and negatively correlated with translation efficiency.

Since our data indicates that G3BP1 targets often possess longer half-life, we hypothesized that G3BP1 may directly stabilize its RNA substrates (Fig. 4a). To further characterize the potential role of G3BP1 in transcript stability, we performed RNA-seq expression analysis in G3BP1/G3BP2 double knockout (G3BP KO) U2OS cells compared against parent U2OS cells (WT) and G3BP KO cells made to express G3BP1 (“rescue”). We observed minimal difference in transcriptome-wide RNA levels between WT and KO cells (Fig. 4b, 4d) and between KO and rescue cells (Fig. 4c, 4e), with median fold-change across all transcripts close to 1 in both comparisons. Interestingly, the 287 G3BP1 targets found in all three TRIBE-ID time points showed statistically significant upregulation in WT cells as compared to KO (Fig. 4b, 4d, 4f). Further, we observed greater stabilization in rescue cells with 80% of G3BP1 TRIBE-ID target transcripts displaying increased abundance compared to KO (Fig. 4c, 4e) and a 1.2-fold increase in median transcript abundance. GO-term analysis of all upregulated transcripts in rescue cells showed similar enriched functions as G3BP1 substrates (Supplementary Fig. 10). Further, most of the 287 G3BP1 substrates showed similar change in abundance in WT and rescue cells as compared to KO cells (Fig. 4h). Overall, our data show that G3BP1 positively regulates RNA stability of its target transcripts in a dose-dependent fashion.

**Figure 4.**
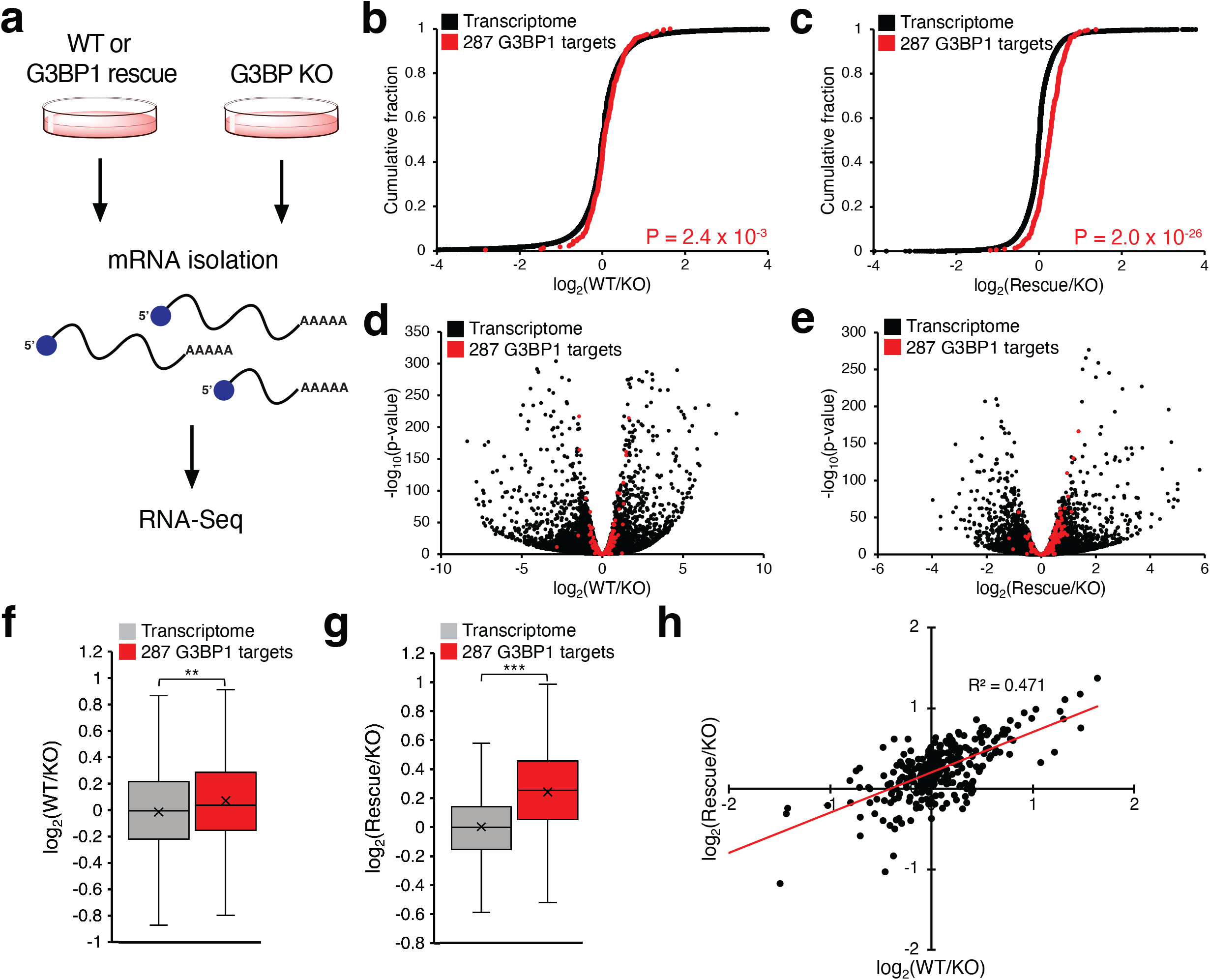
G3BP1 stabilizes target transcripts. **(a)** RNA-seq workflow for analysis of transcript abundance in G3BP KO, WT, and G3BP1 rescue. **(b)** Cumulative distribution of log_2_(fold-change; WT versus G3BP KO) values of all sequenced transcripts and 287 G3BP1 targets identified across three time points with TRIBE-ID. **(c)** Cumulative distribution of log2(fold-change; G3BP1 rescue versus G3BP KO) values of all sequenced transcripts and 287 G3BP1 targets identified with TRIBE-ID. **(d)** Volcano plot of transcript enrichment between WT and KO. **(c)** Volcano plot of transcript enrichment between G3BP1 rescue and KO. **(f)** Boxplot depicting the fold change of transcripts between WT and KO. “x” indicates the mean, and the middle line indicates the median. **(g)** Boxplot depicting the fold change of transcripts between G3BP1 rescue and KO. “x” indicates the mean, and the middle line indicates the median. **(h)** Scatter plot of log_2_(fold-change) values of G3BP1 TRIBE-ID targets in WT and G3BP1-rescue compared against KO. *: p < 0.05; **: p < 0.005; ***: p < 0.0005

### G3BP1-RNA interactions during stress granule formation

The ability to restrict RNA editing to a fixed time window enables the interrogation of RNA-RBP association events as a function of different cell states. We used this property of TRIBE-ID to investigate G3BP1-RNA interactions during stress granule formation. Previously, Khong *et al*.^36^ isolated stress granule cores using sedimentation and G3BP1 affinity enrichment, providing the first stress granule transcriptome data^36^. However, this method does not capture the dynamic outer shell of stress granules which dissociates upon affinity purification and also does not specifically interrogate G3BP1 substrates. More recently, Somasekharan *et al*.^37^ fused APEX2 to G3BP1 in order to capture stress granule transcripts by proximity labeling. Interestingly, the similarities between Khong *et al*.^36^ and Somasekharan *et al*.^37^ are low, leaving questions regarding the composition of stress granule-localized transcripts. In addition, neither of these studies explicitly compared G3BP1-RNA interactions in the cytosol with those in stress granules.

To probe stress granule-associated G3BP1-RNA interactions, we simultaneously treated cells with sodium arsenite to induce stress granules and rapamycin to initiate ADAR editing on G3BP1 substrates (Fig. 5a). Rapamycin-induced editing during sodium arsenite stress resulted in an average of ~3,500 edit sites at 2 hr and ~6,000 edit sites at 4 hr, which is 2-3-fold more than was detected at the corresponding time points in the absence of stress (Fig. 5b and Fig. 3b). On the transcript level, we detected an average of ~1200 edited transcripts at 2 hr and ~1700 edited transcripts at 4 hr, roughly 2-fold more than without arsenite stress (Fig. 5c and 3c). After restricting our analysis to edited transcripts found in both biological replicates, we identified 849 and 1,286 G3BP1 RNA targets at 2 hr and 4 hr of NaAsO_2_ stress, respectively. Our data indicate that localization of G3BP1 in stress granules correlates with an increase in G3BP1-RNA binding, in line with prevailing models proposing a scaffolding role for RNA in templating G3BP1 LLPS^9–11^.

**Figure 5.**
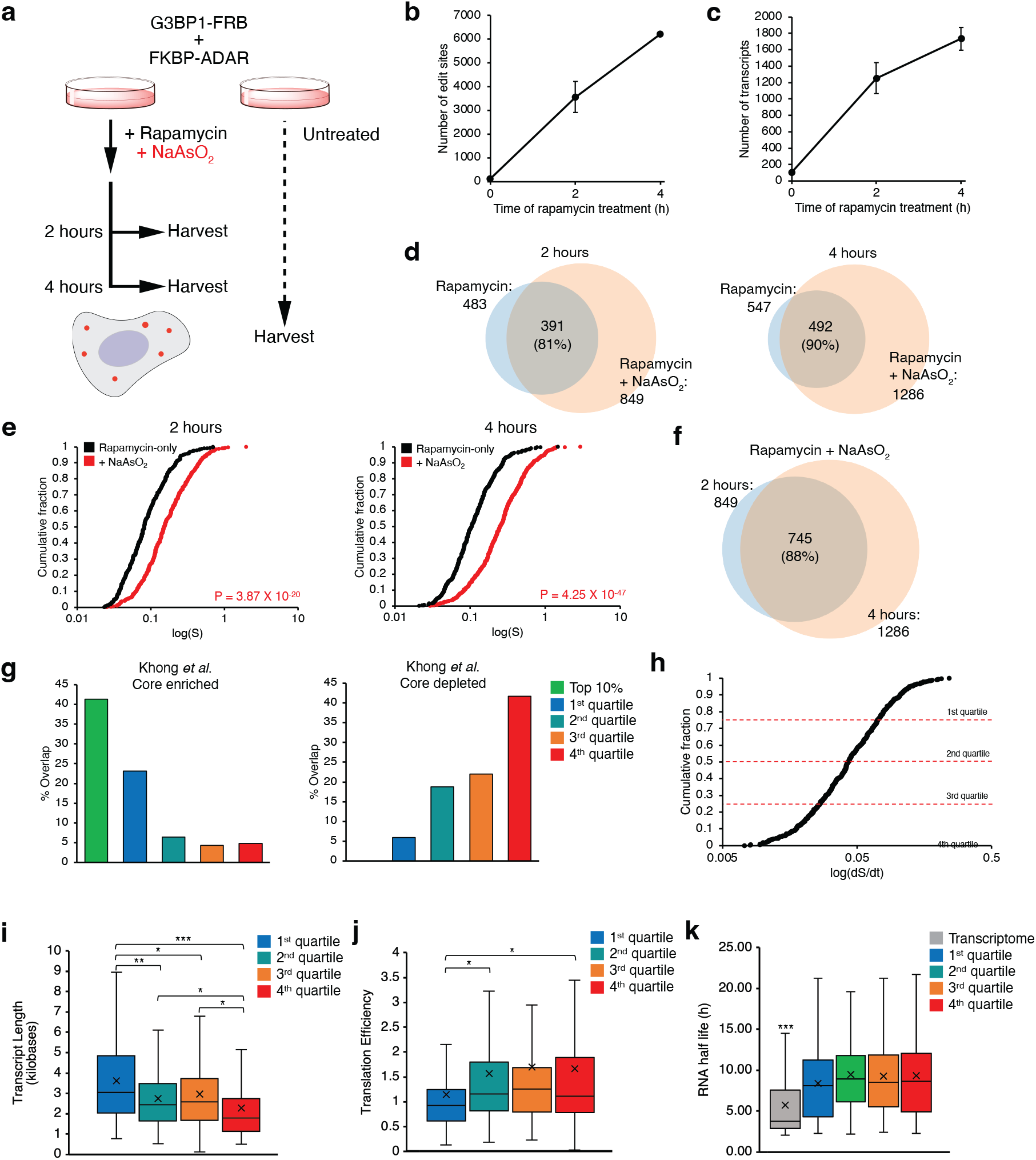
TRIBE-ID captures dynamic G3BP1-RNA interactions during arsenite stress. **(a)** Workflow of G3BP1 TRIBE-ID under NaAsO_2_ stress. **(b)** Average number of edit sites detected with 100 nM rapamycin and 200 μM NaAsO_2_ treatment at each time point. **(c)** Average number of edited transcripts detected with 100 nM rapamycin and 200 μM NaAsO_2_ treatment at each time point. **(d)** Venn diagrams of transcripts found using TRIBE-ID under normal conditions and with NaAsO_2_ treatment at 2 hours and 4 hours. **(e)** Cumulative distribution of S values for TRIBE-ID transcripts detected during normal conditions and with NaAsO_2_ stress at 2 hours and 4 hours. **(f**) Venn diagram of TRIBE-ID transcripts found with NaAsO_2_ treatment at both time points. **(g)** Bar graphs depicting overlap between TRIBE-ID detected transcripts during NaAsO_2_ stress sorted by dS/dt and the stress granule transcriptome data from Khong *et al*. **(h)** Cumulative distribution of NaAsO_2_ exclusive TRIBE-ID transcripts ranked by their dS/dt values. **(i)-(k)** Boxplots depicting the transcript length **(j)**, translation efficiency **(k)**, and half-life **(l)** of transcripts in different dS/dt quartiles of NaAsO_2_ exclusive transcripts. “x” indicates the mean, and the middle line indicates the median. *: p < 0.05; **: p < 0.005; ***: p < 0.0005

We found that the majority of G3BP1-RNA interactions present in unstressed cells were also detected during NaAsO_2_ treatment (Fig. 5d). While this could be explained by detection of residual G3BP1-RNA interactions that remain excluded from stress granules, an alternative hypothesis is that G3BP1-RNA complexes survive stress granule formation. To differentiate between these two scenarios, we compared the amount of editing present on G3BP1 substrates identified in both unstressed and NaAsO_2_ stress conditions using the S parameter defined previously (Fig. 5e). We observed more edits on these transcripts under arsenite stress than without arsenite stress, supporting a model in which G3BP1 recruits its target transcripts to stress granules where they participate in multivalent G3BP1-RNA interactions. Specifically, 85-89% of transcripts found in both stressed and non-stressed conditions at 2 hr or 4 hr had higher S under arsenite stress, and the median S value for G3BP1 transcripts identified with TRIBE-ID during arsenite stress was roughly 2-fold higher than for transcripts from unstressed cells (Fig. 5e). Next, we found 745 transcripts identified with TRIBE-ID shared between 2 hr and 4 hr NaAsO_2_ treatment (Fig. 5f) and sorted them by dS/dt as described previously (Supplementary Fig. 11). We observed that 85% of G3BP1 transcripts found in the top quartile based upon dS/dt were shared between stressed and unstressed cells (Supplementary Fig. 12). Taken together, our data suggests that the vast majority of pre-existing G3BP1-RNA interactions persist during LLPS and are accompanied by the formation of new G3BP1-RNA interactions occurring within stress granules.

Next, we compared our TRIBE-ID G3BP1 stress granule interactome against previously reported stress granule transcriptomic data. Interestingly, we found that transcripts in our dataset with high dS/dt showed substantially higher overlap with the reported stress granule core transcriptome^36^ than those with low dS/dt (Fig. 5g). Inversely, transcripts with low dS/dt overlapped strongly with transcripts that are not reported to partition into stress granule core structures^36^. Our data suggests that TRIBE-ID can identify high affinity G3BP1-RNA interactions and that these RNP complexes tend to partition into stable stress granule core structures as opposed to the dynamic outer shell.

TRIBE-ID analysis of G3BP1-RNA interactions induced by arsenite stress suggested that stress granule formation coincided with a 2.5-fold increase in the number of G3BP1-RNA substrates. To focus on new interactions acquired during stress, we analyzed 481 stress-dependent G3BP1 targets that were not found during TRIBE-ID analysis in unstressed cells. GO-term enrichment analysis of these new transcripts revealed enrichment of RNA metabolism, cell cycle, and translation, which are similar to those found for G3BP1 substrates in the absence of stress (Supplementary Fig. 13). Next, we sorted stress-specific G3BP1 transcripts by dS/dt and grouped them into quartiles as described above (Fig. 5h). Interestingly, we found similar correlations between dS/dt and transcript length, translational efficiency, and half-life as observed for stress-independent G3BP1 substrates (Fig. 5i-k), although the translation efficiency and half-life data were measured in unstressed cells. Together, our data indicates that G3BP1 binds RNA substrates with similar properties with and without stress and suggests that G3BP1 interaction, rather than stress granule localization, may determine the characteristics of transcripts found in stress granules.

### TRIBE-ID detects small molecule-mediated inhibition of G3BP1-RNA interactions

As another application of the TRIBE-ID method, we envisioned that our approach could be applied to detect and characterize direct inhibitors of RNA-protein interactions in live cells. Although small molecules that bind to RNA or RBPs have been identified, their effects on global RNA-protein interactions in living cells remain difficult to measure. As a proof of concept, we chose three reported inhibitors of G3BP1 or G3BP1-RNA interactions: pyridostatin (PDS), resveratrol (RSVL), and epigallocatechin gallate (EGCG). PDS binds and stabilizes G-quadruplexes in RNA and DNA^38,39^ and was reported to inhibit G3BP1-RNA interactions at RNA-G-quadruplexes (rG4)^40^. RSVL and EGCG are reported to bind and inhibit G3BP1 signaling^41–44^, but their effects on G3BP1-RNA interactions have not been described. For this effort, cells were first treated with each compound, and rapamycin was added subsequently to enable editing of RNA targets that remained bound to G3BP1 (Fig. 6a). We observed ~25% reduction in the number of edit sites upon PDS treatment and ~50% reduction with RSVL or EGCG compared to ~8600 edit sites in the rapamycin control (Fig. 6b) (edit sites in the rapamycin control were higher than in previous experiments since samples were sequenced at greater depth). Similarly, we detected an average of ~3200 edited transcripts in control samples and ~2100-2600 edited transcripts from inhibitor treated cells (Fig. 6c). After restricting our analysis to edited transcripts found in both biological replicates, we found 939, 1142, and 1191 of 2108 transcripts edited under rapamycin were lost in the presence of PDS, RSVL, and EGCG, respectively (Fig. 6d), indicating these G3BP1-RNA interactions were completely blocked by small molecule inhibitors. For the transcripts that remained after inhibitor treatments, the global median S values were decreased by 15-30% under all drug treatments compared to rapamycin-only control (Fig. 6e), indicating site-specific or incomplete inhibition of binding to these transcripts by inhibitors. Further, we found 66.4% of transcripts detected in rapamycin-treated cells are reported to contain rG4^38^, and most of the these transcripts were not detected or showed lower editing with PDS treatment (Supplementary Fig. 14), suggesting that PDS inhibits G3BP1 interactions with rG4-containing transcripts. To ensure that the inhibitors do not disrupt components of the TRIBE-ID system other than G3BP1-RNA binding, we confirmed that these molecules do not inhibit ADAR activity, interfere with FRB-FKBP dimerization, or decrease G3BP1 abundance (Supplementary Fig. 15-17).

**Figure 6.**
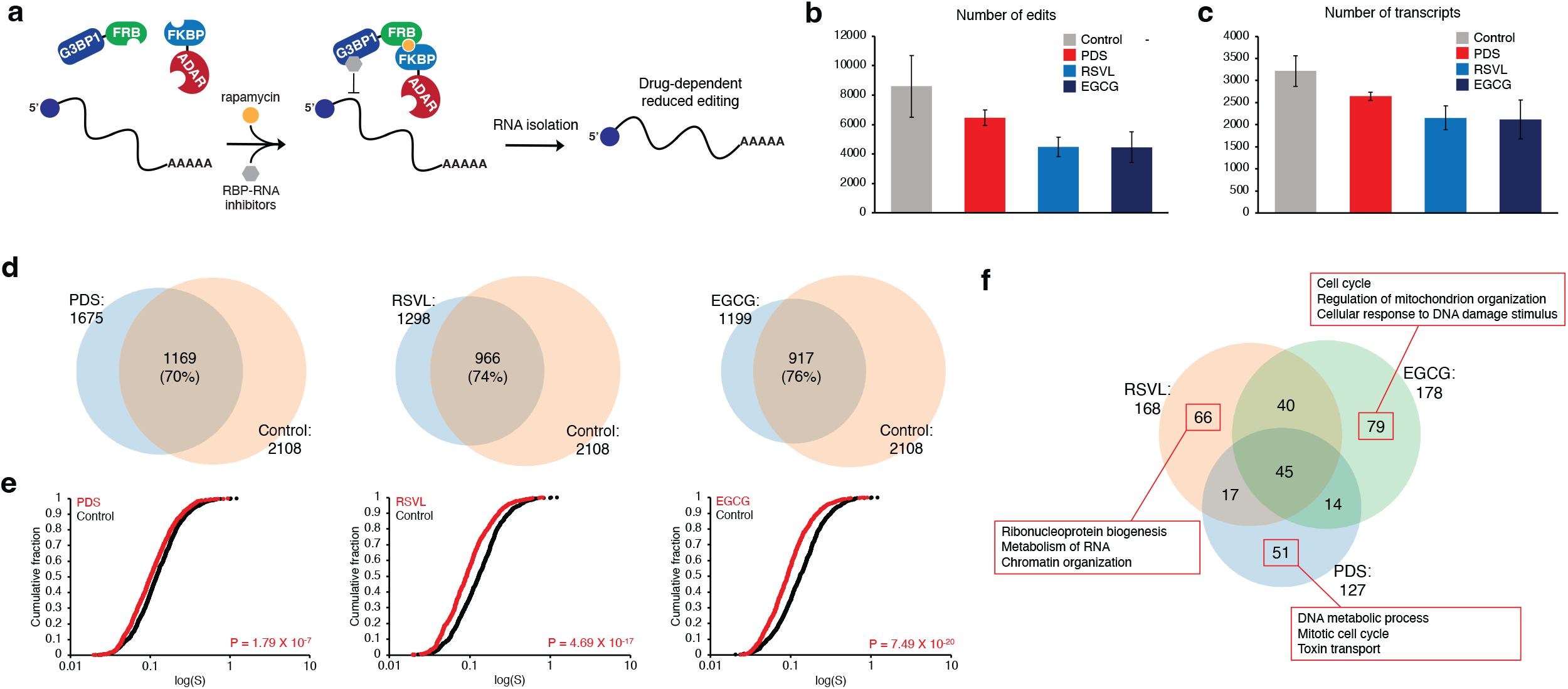
Profiling RBP-RNA drug targets in live cells using TRIBE-ID. **(a)** Workflow for evaluating small molecule G3BP1 inhibitors using TRIBE-ID. **(b)** Average number of TRIBE-ID edit sites detected with inhibitor treatment. **(c)** Average number of TRIBE-ID edited transcripts detected with inhibitor treatment. **(d)** Venn diagram showing overlap between transcripts edited in control (rapamycin only) and transcripts edited with 20 μM PDS, RSVL, or EGCG treatment. **(e)** Cumulative distribution of S values for transcripts detected in control conditions and drug treated conditions. **(f)** Venn diagram showing transcripts with decreased editing under each drug treatment. Top three hits from GO-term enrichment analysis for G3BP1-RNA interactions inhibited upon drug treatment are displayed in red boxes.

Next, we sought to characterize transcripts with significantly reduced G3BP1-RNA interactions in each drug treatment. We selected transcripts that were absent in drug treated cells or those showing a 50% or greater reduction in editing (based upon S value). We found 127, 168, and 178 transcripts such transcripts in PDS, RSVL, and EGCG treated cells, respectively (Fig. 6f). 45 transcripts were shared across all three groups, but 51 transcripts in PDS, 66 in RSVL, and 79 in EGCG were unique in each group. The GO-term enrichment analysis of exclusive transcripts in each group showed enrichment of different cellular processes, ranging from metabolism to cell cycle (Fig. 6f, Supplementary Fig. 18). Taken together, our results demonstrate that TRIBE-ID can detect inhibition of specific G3BP1-RNA interactions due to drug treatment.

## DISCUSSION

In this manuscript, we develop TRIBE-ID, a quantitative method to characterize dynamic and state-specific RNA-protein interactions in live cells. Our approach builds on the TRIBE platform, which provides a facile method to profile RNA-protein binding events in their native context but is not amenable to studying stress-specific interactions or measuring interaction persistence. We apply TRIBE-ID to profile G3BP1 substrates during normal conditions and upon phase separation into stress granules, and also demonstrate its utility in characterizing small molecule inhibitors of RNA-protein binding. Further, we show that most RNA clients are stabilized by G3BP1, and establish correlations between G3BP1 binding, transcript length, and translational efficiency.

RNA-protein condensation has emerged as a ubiquitous biological phenomenon, but the functional role of biomolecular phase separation has remained elusive in many contexts. A key question in understanding this process is to determine the effect of condensation on native RBP-RNA interactions and to elucidate the properties of phase-separated RNPs. Our study profiles the G3BP1-RNA interactome under normal conditions and during NaAsO_2_-induced stress granule formation. We find that high-persistence G3BP1-RNA interactions are preserved and enhanced upon phase separation concomitant with a global increase in G3BP1-bound transcripts and G3BP1-RNA binding events. Interestingly, high-persistence G3BP1-RNA interactions identified with TRIBE-ID overlap strongly with the stress granule core transcriptome identified by Parker and co-workers^36^. We propose a model whereby cytosolic G3BP1-RNA complexes are poised for phase separation and seed stress granule formation as a result of an increase in free RNA concentration (due to decreased ribosomal translation)^45^ and protein oligomerization propensity^9–11^. In support of our model, a recent study demonstrated that artificial tethering of an RNA transcript to G3BP1 resulted in its accumulation in stress granules^46^. Upon condensation, the hierarchy of G3BP1-RNA interactions is preserved with stable interactions predominating at the core and more dynamic RNA-protein interactions found at the periphery, or “outer shell”^47^. New stress-dependent G3BP1-RNA interactions form as a result of RNA recruitment to stress granules via other RBPs, or through stabilization by multivalent G3BP1 binding.

The preservation of the cytosolic G3BP1-RNA interactome upon stress granule formation suggests that the role of condensation may serve to enhance existing G3BP1-RNA interactions. While the effect of G3BP1 binding on mRNA behavior has been in question^6^, our data demonstrates that G3BP1 stabilizes the majority of its substrates in a dose-dependent manner. Further, G3BP1-associated mRNAs exhibit low rates of protein production indicating that an important function of G3BP1 is to stabilize poorly translated mRNAs in order to prevent their degradation. This observation has been made for other stress granule RBPs including DDX1, YB-1, etc.^48–50^ In the context of stress granule condensates, we propose that the high local concentration of G3BP1 protects stress granule-localized transcripts from degradation during the oxidative stress response. In this manner, stress granules can serve as depots to store and protect specific mRNAs during stress when RNA turnover is accelerated^51^; once stress has dissipated, these same mRNAs can be released into the cytoplasm where they can progress through their normal lifecycle.

Beyond profiling native RNA-protein interactions, we demonstrate that TRIBE-ID is a powerful platform for characterizing small molecule RBP inhibitors transcriptome-wide, as temporal control of editing allow us to more reliably distinguish direct from indirect effects of these compounds. Two polyphenol natural products, RSVL and EGCG, have been identified as direct binders of different domains of G3BP1^43,44^, and a recent study showed that PDS abrogated G3BP1 binding to rG4-containing transcripts^40^. Here, we demonstrate that all three compounds inhibit G3BP1-RNA interactions in cells, with RSVL and EGCG showing larger global effects. Since neither RSVL nor EGCG are thought to interact with the G3BP1 RRM domain^43,44^, we propose that these compounds act as allosteric modulators of G3BP1-RNA binding, potentially through modulation of G3BP1 oligomerization. In particular, EGCG binding to G3BP1 requires the RGG domain^44^, which is important for protein-protein interactions and RNA recognition^9–11^. Given the interest in polyphenols for therapeutic application in cancer, aging, and neurodegeneration, their effect on G3BP1-RNA interactions should also be considered when accounting for mechanism of action. More broadly, our work demonstrates how TRIBE-ID can be utilized as a novel RBP-RNA inhibitor screening platform.

Finally, while the original TRIBE method employed *Drosophila* Adar catalytic domain in fruit flies^22^, we find that human ADAR2 E488Q outperforms the *Drosphila* enzyme in human cells as it exhibits lower background editing and higher on-target editing efficiency when fused to G3BP1. Further, the human E488Q mutant exhibits superior TRIBE editing as compared to human E488Q/T375G double mutant that was developed for single-site editing using the CRISPR/Cas13-based system^30^. Together, our data suggest that selection of adenosine deaminase domain can dramatically affect the outcome of TRIBE and TRIBE-ID experiments, and that examination of different ADAR domains may be valuable when designing TRIBE-based RBP profiling experiments.

In summary, our study offers a novel and general approach to profile RBP-RNA interactions with temporal control. We propose that TRIBE-ID can be applied broadly to profile RNPs and RNA-protein condensates and to measure dynamic RNA-protein interactions on rapid time scales. We also envision that leveraging diverse RNA-modifying enzymes^23,24^ and inducible dimerization tools could further improve our method. Such studies are in progress in our lab and will provide new insights into RNA-protein interactions in biology.

## DATA AVAILABILITY

The sequencing data reported in this paper have been deposited in the NCBI Gene Expression Omnibus (accession code: GSE207005).

## ACKNOWLEDGEMENTS

We thank D.W. Sanders and C. Brangwynne for providing the G3BP1/2 KO cell line. We thank C. DeCoste and K. Rittenbach at the Princeton University Flow Cytometry Resource Facility for assistance with cell sorting experiments. We thank W. Wang and R. Leach at the Princeton University Genomics Core Facility for performing Illumina sequencing and assisting K.W.S. with bioinformatic analysis. R.E.K. acknowledges support from the National Institute of Health (R01 GM132189), the Sidney Kimmel Foundation and the Alfred P. Sloan Foundation. K.W.S. was generously supported by the Edward C. Taylor 3rd Year Graduate Fellowship in Chemistry. All authors thank Princeton University for financial support.

## AUTHOR INFORMATION

Department of Chemistry, Princeton University, Princeton, NJ, USA

Kyung W. Seo & Ralph E. Kleiner

## AUTHOR CONTRIBUTIONS

R.E.K. conceived the study, designed experiments, wrote the manuscript, and supervised K.W.S. K.W.S. performed all experiments and bioinformatic analysis and wrote the manuscript.

## METHODS

### Plasmids

G3BP1 cDNA was obtained from Genscript (OHu02150D). FRB, FKBP-GFP, dADAR(E488Q), and hADAR2 WT cDNA were obtained from Addgene (#104476, 106924, 154786, 103866) The hADAR2 E488Q and T375G mutations were introduced into hADAR using overlap extension PCR with mutagenic primers. Fusion proteins were assembled with overlap extension PCR. For transient transfection and construction of cell lines, cDNAs were cloned into pcDNA5/FRT/TO vector (Life Technologies, V6520–20) or pBABE-puro vector (Addgene, #1764).

### Cell culture

All mammalian cells were cultured at 37°C in a humidified atmosphere with 5% CO2 in DMEM (Thermo Fisher, 11995073) supplemented with 10% fetal bovine serum (Bio-Techne, S12450H), 1x penicillin-streptomycin (Thermo Fisher 15070-063) and 2 mM L-glutamine (Thermo Fisher, 25030–081).

### Generation of stable cell lines

To generate a stable cell line expressing G3BP1-FRB or 3xFLAG-G3BP1, the Flp-In T-Rex 293 cells were seeded at 0.4 x 10^6^ cells per well in a six-well plate. Next day, the cells were co-transfected using Lipofectamine 2000 (Thermo Scientific, 11668027) with pOG44 (2 μg; Thermo Fisher, V600520) and pcDNA5/FRT/TO plasmid containing the gene of interest (0.2 μg). Colonies were formed after selection in 100 μg/mL hygromycin B and 15 μg/mL blasticidin. To confirm expression of the proteins, cells were grown in the presence or absence of 1 μg/mL tetracycline for 24 hours and lysed with NP-40 buffer (50 mM Tris-Cl, pH 7.5, 150 mM NaCl, 1% NP-40) supplemented with 1 mM phenylmethyl sulfonyl fluoride (PMSF) and protease inhibitor tablet (Sigma, 11836170001). The proteins were separated on SDS-PAGE gels and analyzed with western blot (anti-G3BP1, 1:1,000 dilution, Santa Cruz, H-10; anti-FLAG M2, 1:1,000 dilution, Sigma, F1804).

To generate a stable cell line expressing both G3BP1-FRB and FKBP-hADAR(E488Q), AmphoPack-293 cells were seeded at 2.0 x 10^6^ per 10-cm plate on day 1. 24 hours later, the cells were transfected using Lipofectamine 2000 with pBABE-puro vector containing FKBP-hADAR(E488Q)-GFP gene (20 μg). On day 3, transfection medium was removed, and fresh medium was added. On day 3, Flp-In T-Rex 293 cells expressing G3BP1-FRB were also seeded at 2.0 x 10^6^ per 10-cm plate. Medium containing virus was collected three times on days 4 and 5, filtered through a 0.45 μm filter, and used immediately for infection. The virus-containing medium was supplemented with 2 μg/mL Polybrene and directly added to Flp-In T-Rex 293 cells. Colonies were formed after selection in 1 μg/mL puromycin and sorted using FACSAria Fusion cytometer. After recovery, the sorted cells were harvested after 1 μg/mL tetracycline induction, and the expression of both G3BP1-FRB and FKBP-hADAR(E488Q) proteins was confirmed with western blot (anti-G3BP1, 1:1,000 dilution, Santa Cruz, H-10; anti-GFP, 1:2,000 dilution, Abcam, ab290).

### Immunofluorescence microscopy

Cells stably expressing G3BP1-FRB and FKBP-hADAR(E488Q) were seeded in a 6-well plate with 12-mm glass coverslips. The cells were treated with 100 nM rapamycin with or without 200 μM NaAsO_2_ for 2 hours and fixed with 3% paraformaldehyde in PBS for 15 min at room temperature and washed with PBS twice. The fixed cells were permeabilized with PBST (0.1% Triton X-100) for 15 min at room temperature and blocked with 5% goat serum in PBST for 30 min at room temperature. The cells were incubated with anti-G3BP1 antibody (1:200 dilution, Santa Cruz, H-10) and anti-GFP antibody (1:200 dilution, Abcam, ab290) for 2 h at room temperature and washed with PBST (0.1% Tween-20) for 5 min three times. Next, cells were incubated with goat anti-rabbit Alexa 488 antibody (1:800 dilution, Jackson ImmunoResearch, 111-005-144) and goat anti-mouse Alexa 594 antibody (1:800 dilution, Jackson ImmunoResearch, 115-585-003) for 30 min at room temperature. The cells were washed twice with PBST, stained with Hoechst 33342 (1 μg/mL, Thermo Scientific, H3570) in PBST for 5 min, and washed with PBS twice for 5 min. The coverslips were mounted in ProLong Gold AntiFade Mountant (Invitrogen, P36930) and imaged using NIS Elements AR software and a Nikon Eclipse Ti microscope equipped with a ×100 objective and CMOS camera. Images used for direct comparison were acquired using standardized illumination and exposure settings and displayed with identical lookup table settings.

### Transient transfection

For TRIBE analysis, 2 x 10^6^ HEK293T cells were seeded per 10-cm plate 24 hours before transfection. The cells were transfected using Lipofectamine 2000 with 5 μg pcDNA5 vector encoding either ADAR alone or G3BP1-ADAR fusion construct and 5 μg pcDNA5 vector encoding GFP per 10-cm plate. The transfected cells were sorted using FACSAria Fusion cytometer. GFP-positive cells were collected, and their total RNA was isolated with TRIzol (Thermo Fisher, 15596018) following the manufacturer’s protocol. For G3BP KO analysis, G3BP KO U2OS cells were plated and transfected using Lipofectamine 2000 with pcDNA5 vector encoding G3BP1 as described for ADAR constructs above. Total RNA was isolated with TRIzol (Thermo Fisher, 15596018) following the manufacturer’s protocol.

### Immunoprecipitation and RIP-Seq

For co-immunoprecipitation of G3BP1-FRB and FKBP-ADAR in the presence of rapamycin, 2 x 10^6^ Flp-In T-Rex 293 cells that express both G3BP1-FRB and FKBP-ADAR were plated in 10-cm plates. Two 10-cm plates were used per condition per replicate. Next day, the cells were induced with 1 μg/mL tetracycline for 24 hr after which cells were treated with 20 μM PDS for 4 hours, 20 μM RSVL for 3 hours, or 20 μM EGCG for 1 hour. Then, 100 nM rapamycin was added to cells, and they were harvested after 2 hours. The cells were lysed with Buffer A (150 mM KCl, 10 mM HEPES, pH 7.6, 2 mM EDTA, 0.5% NP-40, 0.5 mM DTT) supplemented with 1 mM PMSF and protease inhibitor tablet (Sigma, 11836170001) and centrifuged at 15,000 G for 20 min at 4°C to clear the lysate. To minimize non-specific binding of proteins to the beads, each lysate was incubated with 20 μL Protein-G beads for 1 hour at 4 °C. Anti-G3BP1 beads were generated by incubating 200 μL Protein-G beads with 20 μL anti-G3BP1 antibody (Abcam, ab290) in 1 mL Buffer B (200 mM NaCl, 50 mM HEPES, pH 7.6, 2 mM EDTA, 0.05% NP-40, 0.5 mM DTT) for 1 hour at room temperature. The pre-cleared lysate was incubated with 40 μL anti-G3BP1 beads for 4 hours at 4 °C. The beads were then washed eight times with 1 mL ice cold Buffer B, and proteins were eluted for western blot analysis by boiling the beads in sample buffer.

For RIP-Seq, Flp-In T-Rex 293 cells that express 3xFLAG-G3BP1 were used, and the following changes were made to the protocol described above. All the buffers were supplemented with Ribolock RNase inhibitor (Thermo Fisher, EO0381; 1:100). Anti-FLAG antibody (Sigma, F1804) was used, and the proteins were eluted by incubating the beads with 200 μg/mL 3xFLAG peptide (APExBIO, A6001) for 2 hours at 4 °C. RNA bound to the eluted proteins was isolated with TRIzol LS reagent (Thermo Fisher, 10296010), and the poly(A) RNA was isolated with two rounds of selection using oligo-(dT)_25_ beads (NEB, S1419S). The purified poly(A) RNA was used for RNA-Seq library generation using the NEBNext Ultra II Directional Library Prep kit for Illumina (NEB, E7760S). Amplified cDNAs were submitted for Illumina sequencing.

### RNA-Seq for TRIBE, TRIBE-ID, and G3BP KO

For screening ADARs, we extracted total RNA with TRIzol reagent (Thermo Fisher, 15596018) 24 hours after transfection and immediately after cell sorting by FACS. For TRIBE-ID, the cells expressing G3BP1-FRB and FKBP-hADAR(E488Q) were treated with 100 nM rapamycin and harvested after 2, 4, or 8 hours. In parallel, same cells were treated with 100 nM rapamycin and 200 μM NaAsO_2_ and harvested after 2 or 4 hours. For drug treatment, the cells expressing G3BP1-FRB and FKBP-hADAR(E488Q) were pre-treated with 20 μM pyridostatin (PDS) for 4 hours, 20 μM resveratrol (RSVL) for 3 hours, or 20 μM epigallocatechin gallate (EGCG) for 1 hour. 100 nM rapamycin was added to the pre-treated cells, and they were harvested after 2 hours of rapamycin treatment. For G3BP KO rescue experiment, the KO, parental WT, and rescue cells were harvested. All total RNAs were extracted using TRIzol. Poly(A) RNA was isolated with two rounds of selection using oligo-(dT)25 beads (NEB, S1419S). 50 ng of poly(A) RNA from each sample was used for RNA-Seq library generation using the NEBNext Ultra II Directional Library Prep kit for Illumina (NEB, E7760S). Amplified cDNAs were submitted for Illumina sequencing.

### Bioinformatic analysis

The A-to-I mutation analyses were performed as described in HyperTRIBE^29^. Briefly, sequencing reads were trimmed and mapped to the hg38 using STAR aligner (v.2.7.0f). Only uniquely mapped reads were used for later steps. Nucleotide frequency at each position in the transcriptome was recorded from aligned reads and uploaded to MySQL databases. The ADAR-transfected RNA nucleotide frequency was compared to the wild-type RNA nucleotide frequency to identify RNA edit sites and calculate the percentage of editing at each site. Only mutated sites with >20 read counts were selected for further analyses. For transcript analyses, we eliminated transcripts with only intronic edit sites and kept transcripts detected in both replicates. The S value of each transcript was calculated with this equation:

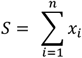

where n is the number of edit sites and x is edit fraction at each site.

Sequences fifty bases upstream and downstream were extracted from the reference and used for motif analysis by DREME (4.12.0). Gene ontology analysis was performed using Metascape software available online (http://metascape.org)^52^. All length data was obtained using Ensemble’s Biomart tool. Translational efficiency values were calculated from Sidrauski *et al*.^34^ (ribosome bound fragment reads / RNA-Seq reads). Half-life data were acquired from Tani *et al*.^33^, and half-lives of >24 h were not considered in the analysis.

For RIP-Seq and G3BP KO analysis, sequencing reads were trimmed and mapped to the GRCh38 using STAR aligner. Uniquely mapped bam files were used for DESeq2 (v.2.11.40.6) to calculate the differential abundance of transcripts. For RIP-Seq, transcripts with positive fold change in IP with adjusted p-value < 0.05 were considered enriched in G3BP1.

