## Supplementary Information for "Profiling dynamic RNA-protein interactions using small molecule-induced RNA editing"

Table of contents

i. Supplementary Table 1
ii. Supplementary Figure 1
iii. Supplementary Figure 2
iv. Supplementary Figure 3
v. Supplementary Figure 4
vi. Supplementary Figure 5
vii. Supplementary Figure 6
viii. Supplementary Figure 7
ix. Supplementary Figure 8
x. Supplementary Figure 9
xi. Supplementary Figure 10
xii. Supplementary Figure 11
xiii. Supplementary Figure 12
xiv. Supplementary Figure 13
xv. Supplementary Figure 14
xvi. Supplementary Figure 15
xvii. Supplementary Figure 16
xviii. Supplementary Figure 17
xix. Supplementary Figure 18

| Samples | Uniquely mapped reads | |
|---|---|---|
| | Rep 1 | Rep 2 |
| (TRIBE experiment) | | |
| Untransfected | $48.3 \times 10^6$ | $60.7 \times 10^6$ |
| dADAR(E488Q) | $16.9 \times 10^6$ | $38.2 \times 10^6$ |
| hADAR(E488Q) | $19.6 \times 10^6$ | $35.6 \times 10^6$ |
| hADAR(E488Q/T375G) | $27.0 \times 10^6$ | $50.4 \times 10^6$ |
| G3BP1-dADAR(E488Q) | $40.9 \times 10^6$ | $61.3 \times 10^6$ |
| G3BP1-hADAR(E488Q) | $28.1 \times 10^6$ | $52.4 \times 10^6$ |
| G3BP1-hADAR(E488Q/T375G) | $22.5 \times 10^6$ | $42.3 \times 10^6$ |
| (TRIBE-ID) | | |
| Untreated | $19.7 \times 10^6$ | $46.2 \times 10^6$ |
| Rapamycin 2 hours | $18.7 \times 10^6$ | $53.4 \times 10^6$ |
| Rapamycin 4 hours | $25.4 \times 10^6$ | $55.8 \times 10^6$ |
| Rapamycin 8 hours | $27.1 \times 10^6$ | $62.6 \times 10^6$ |
| Rapamycin + $NaAsO_2$ 2 hours | $25.4 \times 10^6$ | $28.1 \times 10^6$ |
| Rapamycin + $NaAsO_2$ 4 hours | $20.9 \times 10^6$ | $31.3 \times 10^6$ |
| (Small molecule inhibitors) | | |
| Untreated | $35.0 \times 10^6$ | $48.6 \times 10^6$ |
| Rapamycin | $40.0 \times 10^6$ | $38.2 \times 10^6$ |
| Rapamycin + PDS | $33.0 \times 10^6$ | $45.1 \times 10^6$ |
| Rapamycin + RSVL | $33.5 \times 10^6$ | $41.3 \times 10^6$ |
| Rapamycin + EGCG | $43.4 \times 10^6$ | $36.8 \times 10^6$ |

**Supplementary Table 1**: Number of uniquely mapped reads TRIBE and TRIBE-ID RNA sequencing samples.

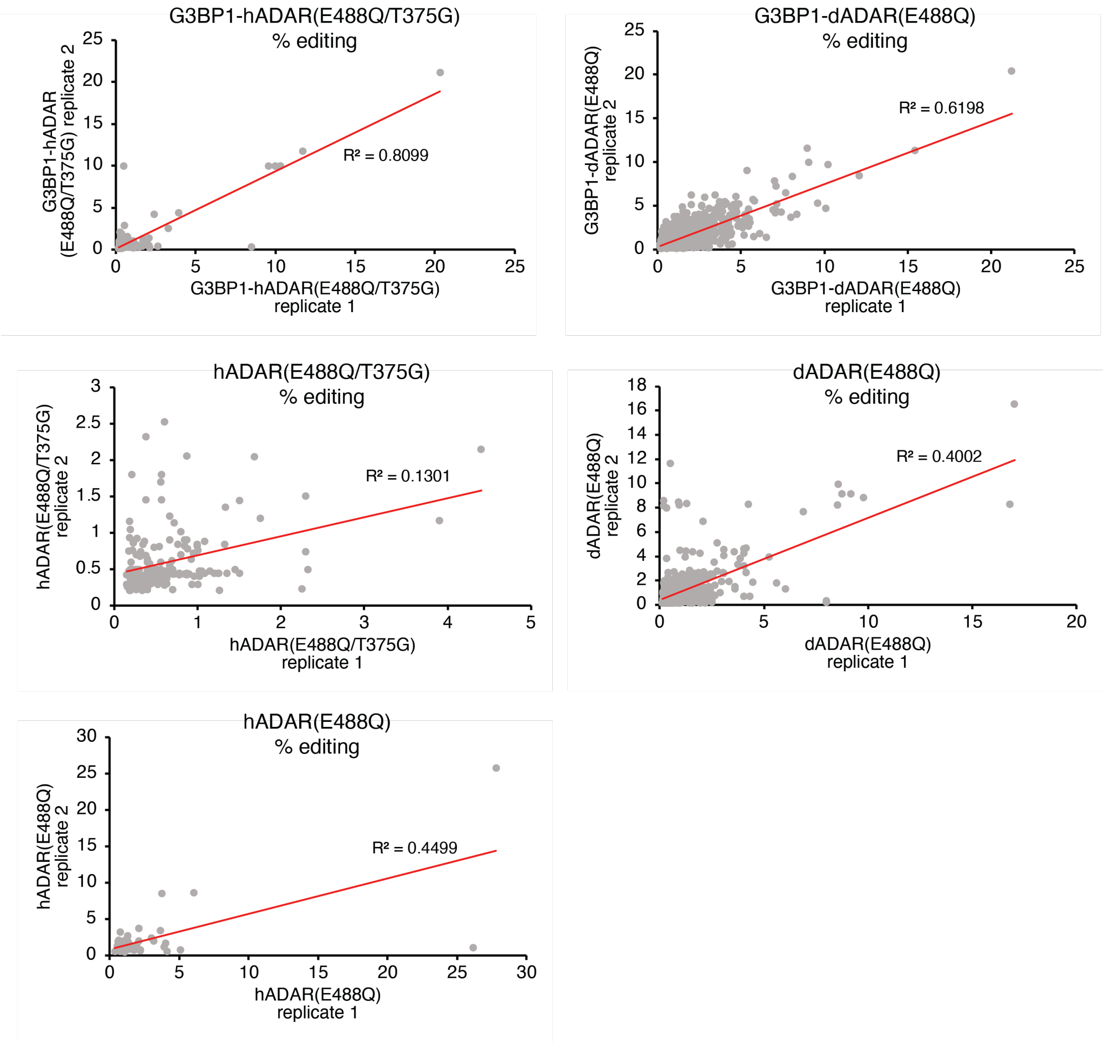


**Supplementary Figure 1**: Reproducibility of editing between replicate 1 and replicate 2 in cells transfected with G3BP1-ADAR or ADAR alone.

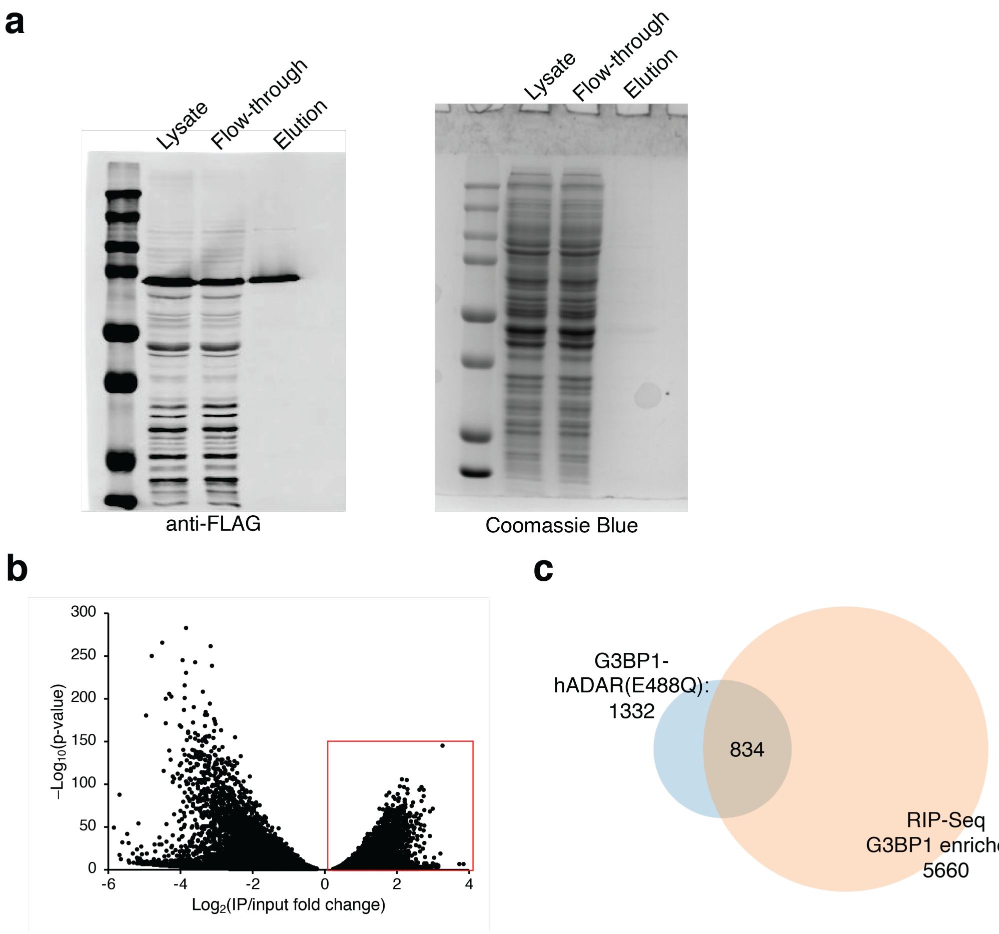


**Supplementary Figure 2**: G3BP1 RIP-seq experiment. **(a)** Western blot analysis FLAG-tagged G3BP1 immunoprecipitation. The same samples were run on a separate gel and stained with Coomassie Blue to visualize all proteins in each sample. Three independent biological replicates were performed. **(b)** Volcano plot showing enriched transcripts in FLAG-G3BP1 RIP-Seq. The transcripts in red-outlined box show statistically significant enrichment ($p$<0.05). **(c)** Venn diagram showing overlap between enriched transcripts in RIP-Seq and transcripts identified in G3BP1-hADAR(E488Q) TRIBE experiment.

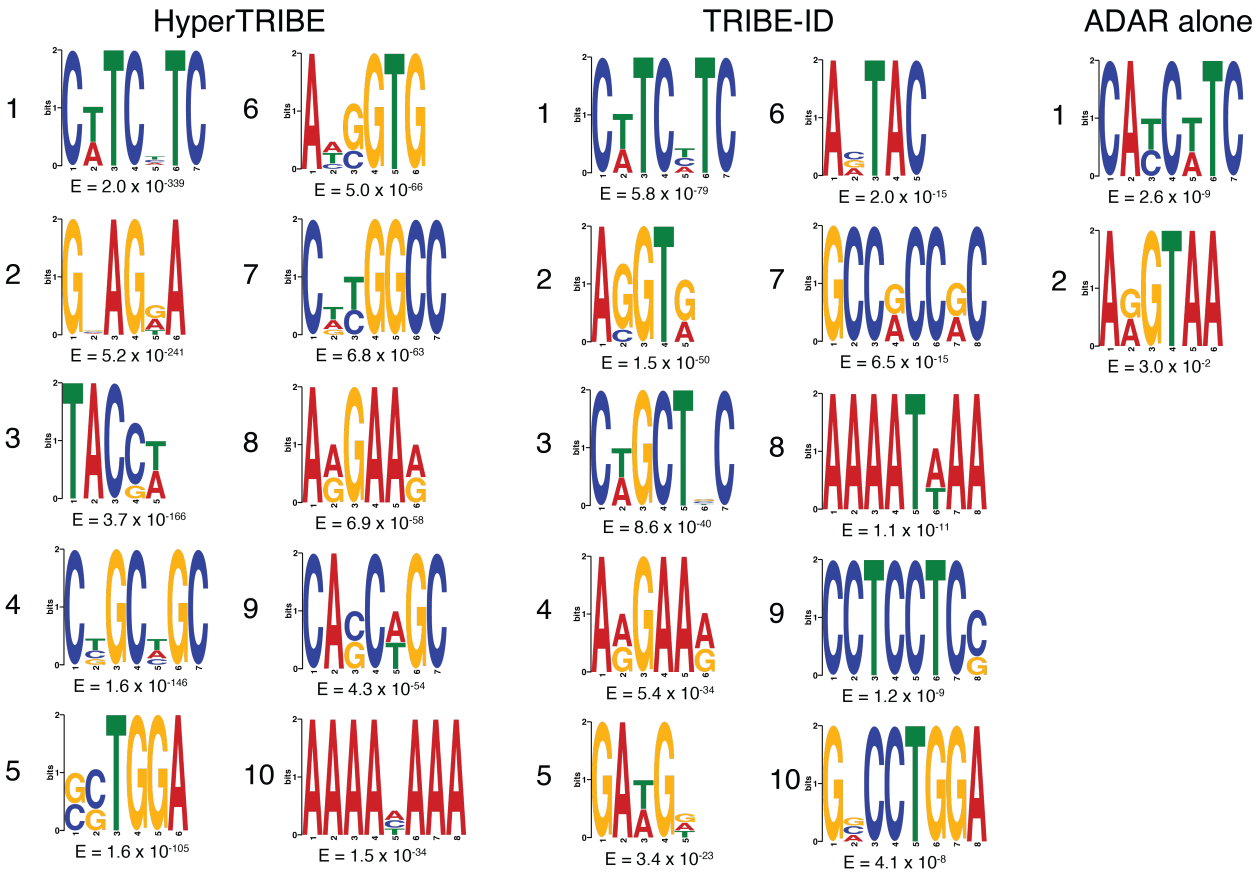


**Supplementary Figure 3**: Top 10 most significant sequence motifs identified with DREME in 200 nt region surrounding edit sites from G3BP1-hADAR(E488Q) (TRIBE experiment), TRIBE-ID (287 transcripts found at 2 hr, 4 hr, and 8 hr), and hADAR(E488Q) (TRIBE control) data.

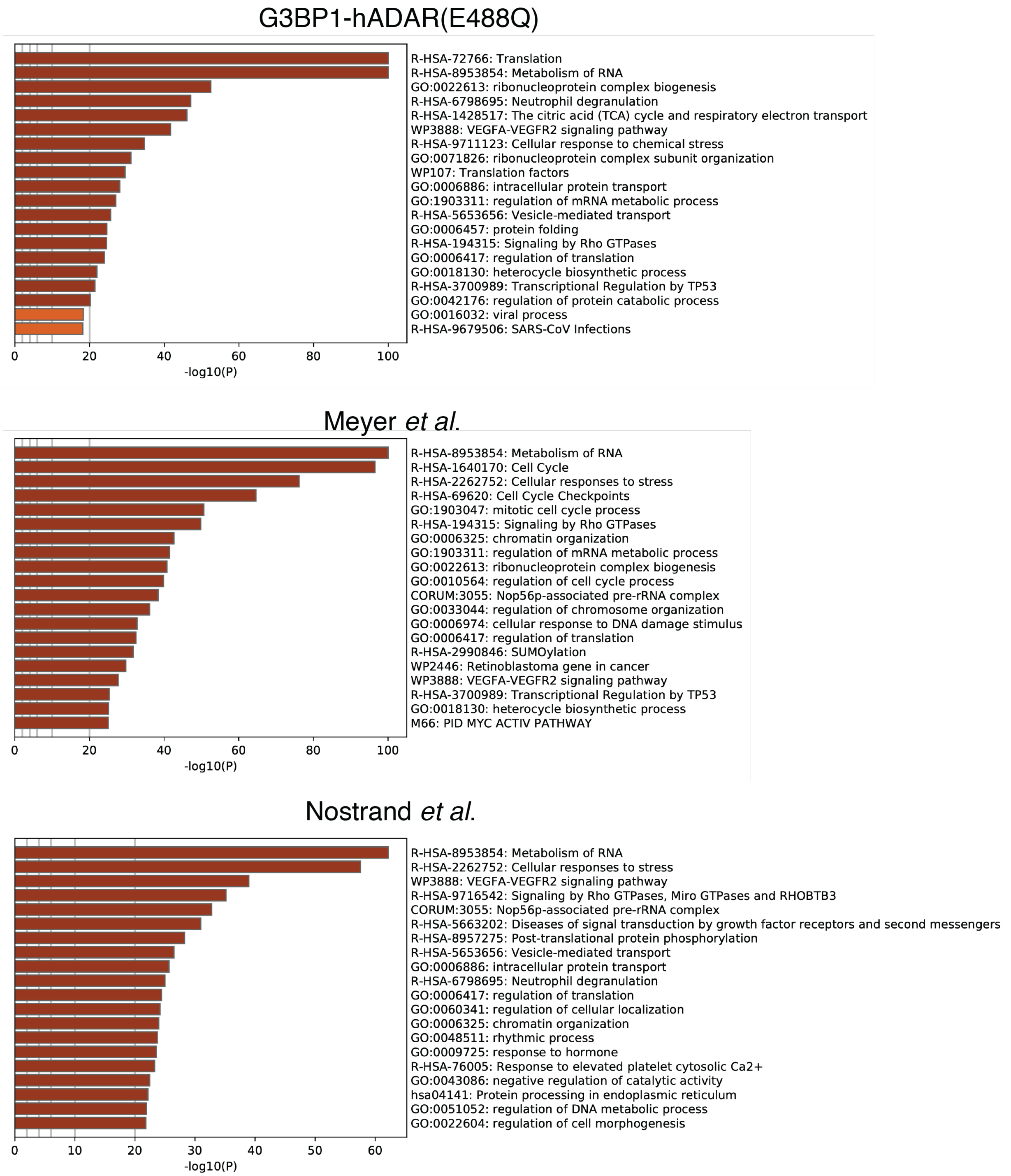


**Supplementary Figure 4**: Top 20 enriched gene ontology terms for G3BP1 target transcripts found in G3BP1 TRIBE experiment, Meyer *et al*.[1] PAR-CLIP, and Van Nostrand *et al*.[2] eCLIP. For CLIP data, top 1000 most abundant transcripts (ranked by number of identified peaks) were subjected to GEO analysis.

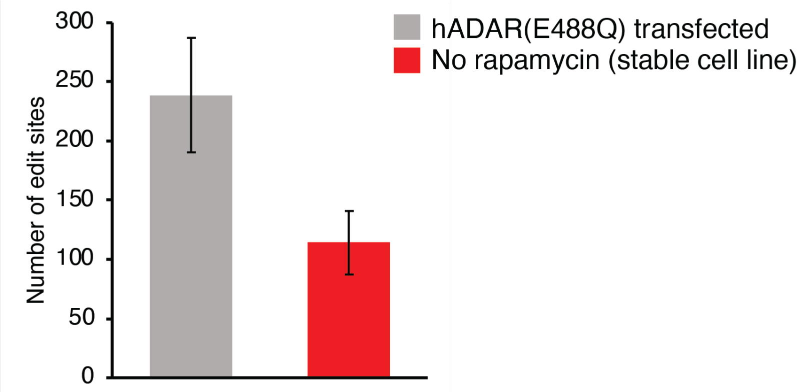


**Supplementary Figure 5**: Background editing from hADAR(E488Q) construct. HEK293T transfected with hADAR(E488Q) or stable HEK293 cell line expressing G3BP1-FRB and FKBP-ADAR without rapamycin treatment were analyzed using TRIBE bioinformatic platform using WT HEK293T as control sample.

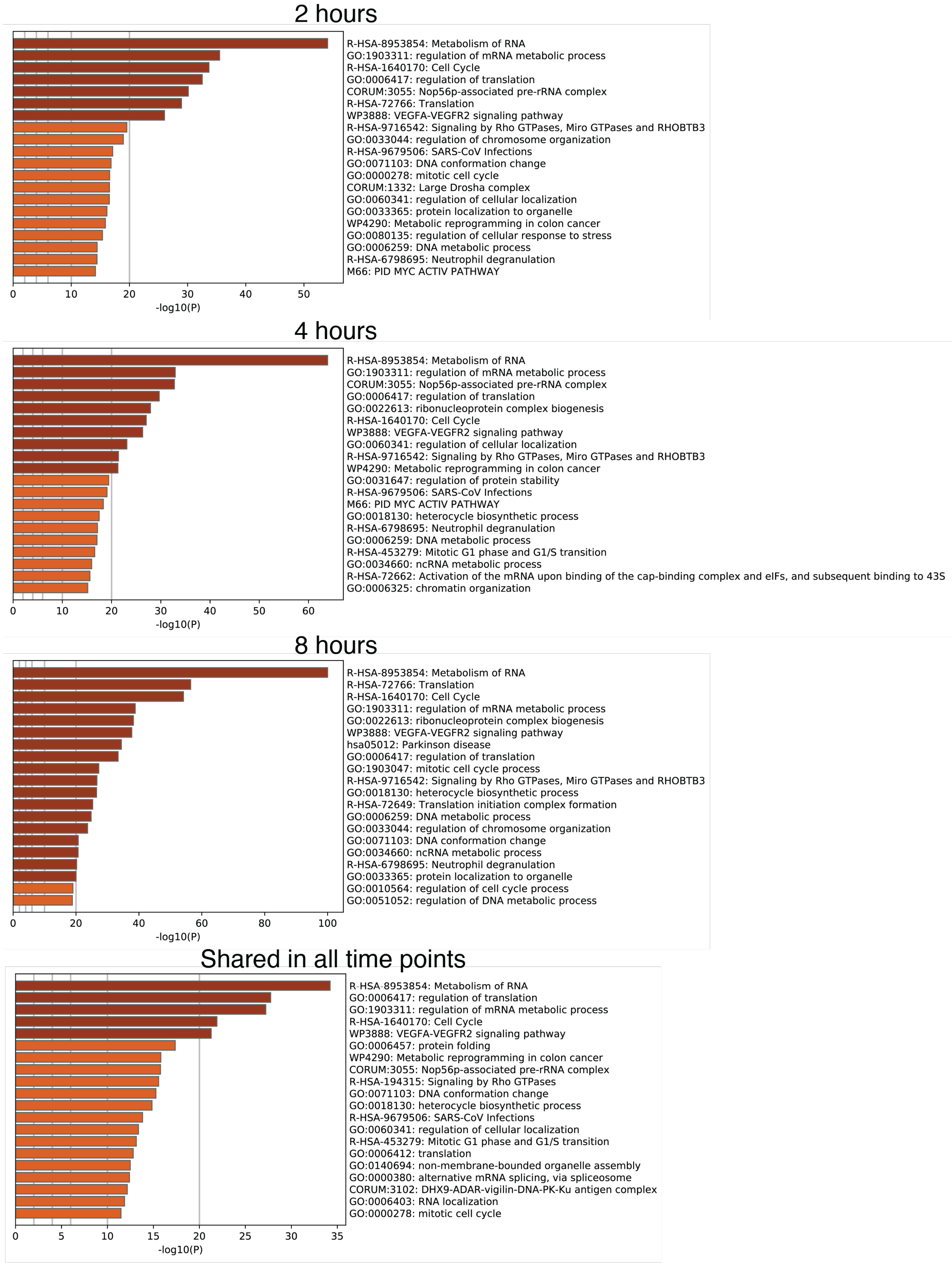


**Supplementary Figure 6**: Top 20 enriched gene ontology terms for G3BP1 TRIBE-ID transcripts detected at 2 hr, 4 hr, 8 hr, or shared across all three time points.

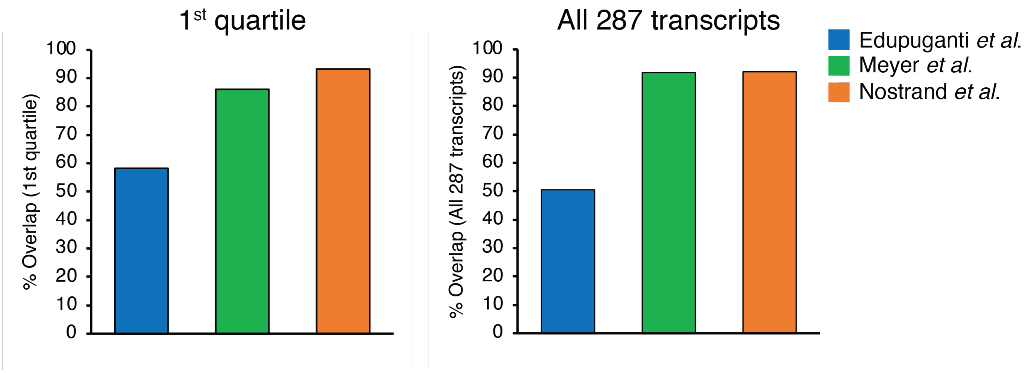


**Supplementary Figure 7**: Bar graphs depicting the overlap between transcripts detected in G3BP1 TRIBE-ID experiment and G3BP1 CLIP data from Edupuganti *et al*.[3], Meyer *et al*.[1], and Van Nostrand *et al*.[2]. 1st quartile and shared 287 transcripts from TRIBE-ID are defined in Fig. 3e and 3g.

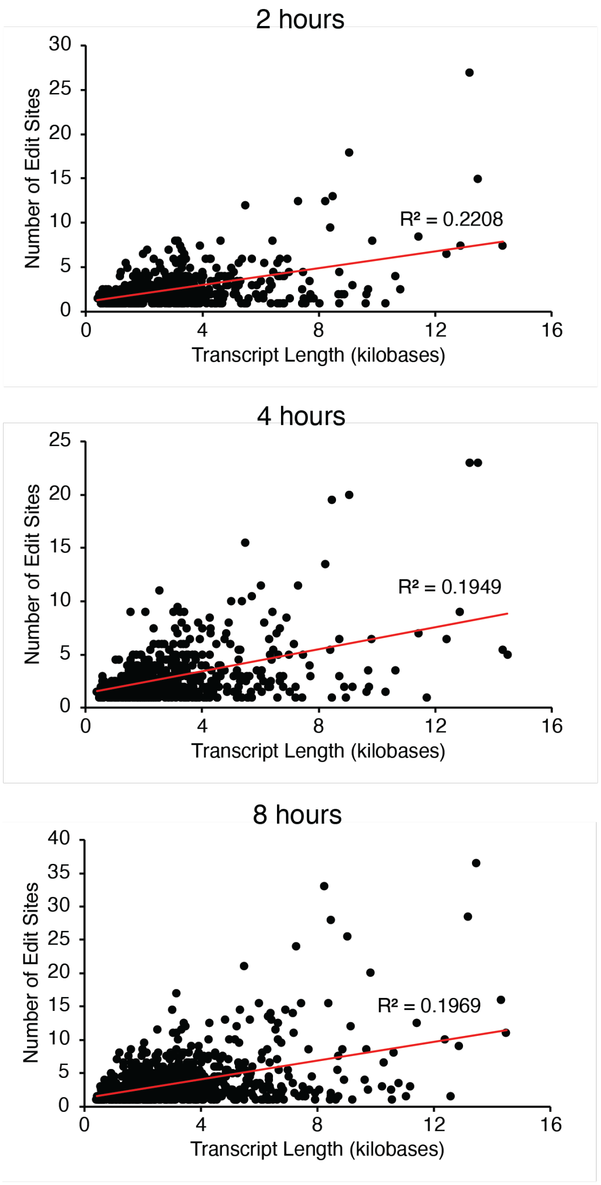


**Supplementary Figure 8**: Scatter plots showing correlation between transcript length and number of edit sites from G3BP1 TRIBE-ID experiment.

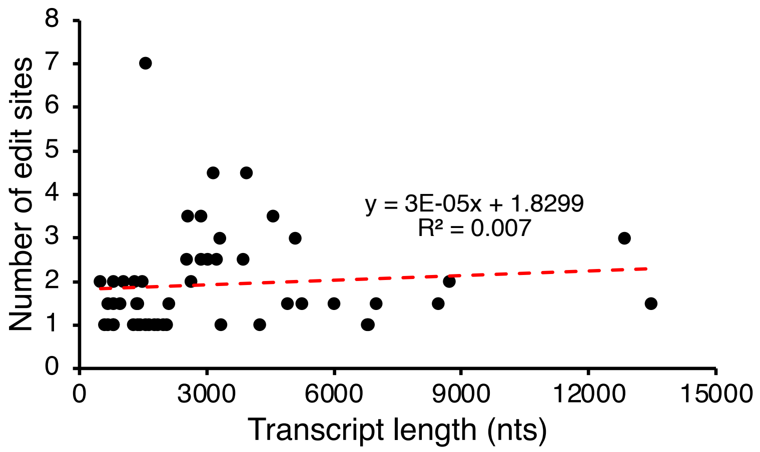


**Supplementary Figure 9**: Scatter plots showing correlation between transcript length and number of edit sites from TRIBE control experiment (hADAR(E488Q)).

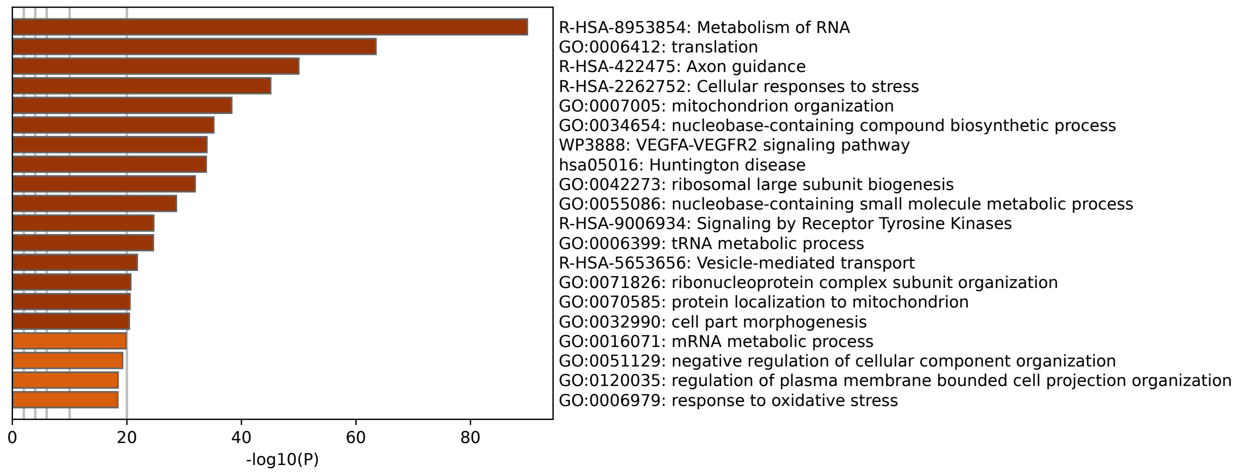


**Supplementary Figure 10**: Top 20 enriched gene ontology terms for transcript upregulated in G3BP1 rescue cells as compared to G3BP KO.

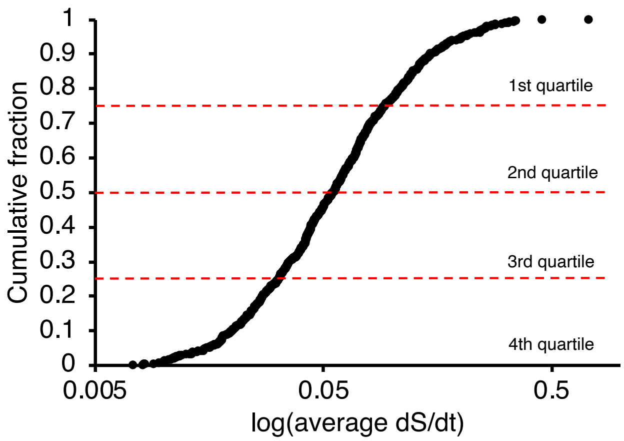


**Supplementary Figure 11**: Cumulative distribution of 745 transcripts detected in G3BP1 TRIBE-ID experiment under $NaAsO_2$ stress, ranked by dS/dt values. Dotted red lines separate quartiles.

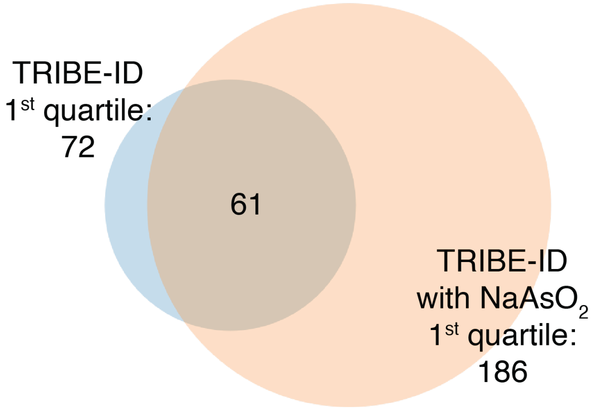


**Supplementary Figure 12**: Venn diagram showing overlap between top quartile transcripts in TRIBE-ID experiment with and without $NaAsO_2$ stress.

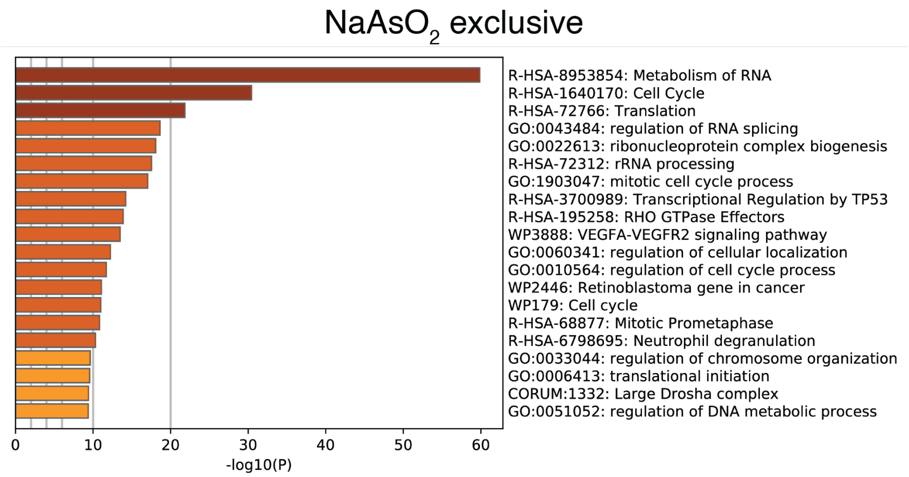


**Supplementary Figure 13**: Top 20 enriched gene ontology terms for 481 G3BP1 TRIBE-ID transcripts detected only under $NaAsO_2$ treatment and detected in unstressed cells.

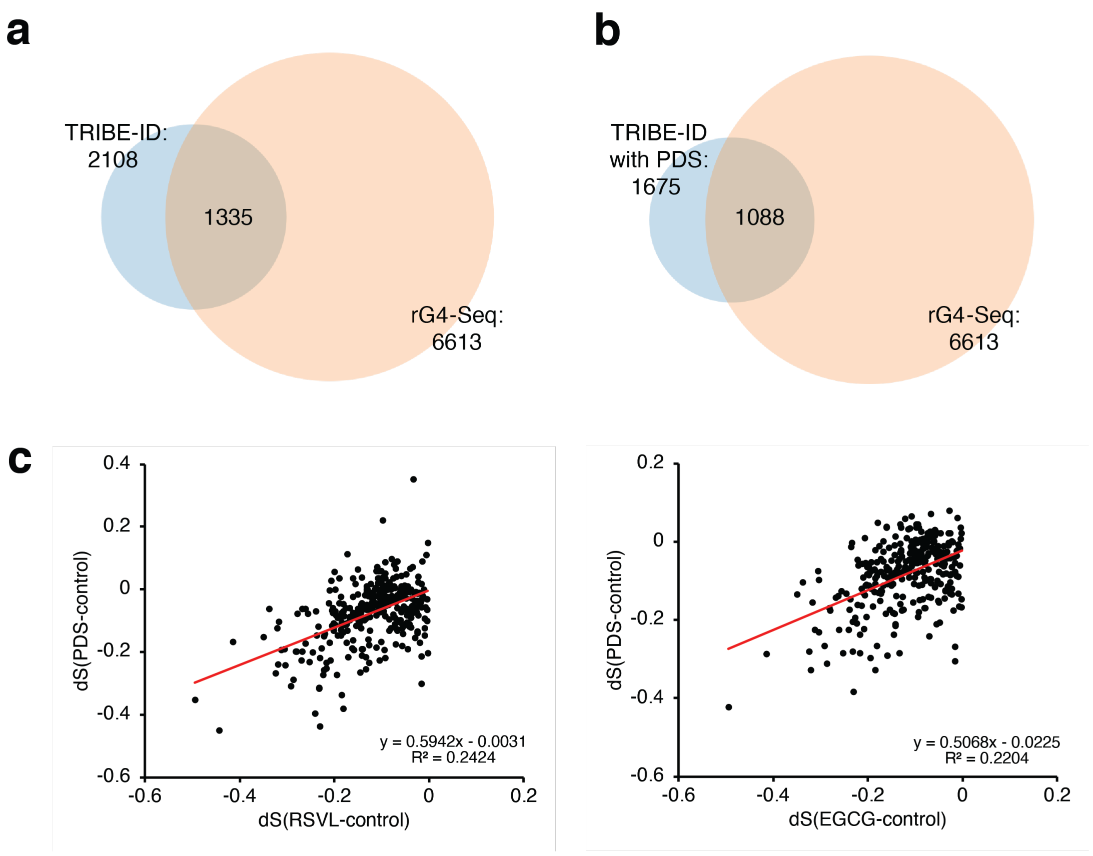


**Supplementary Figure 14**: PDS affects editing on transcripts containing rG4. **(a)** Venn diagram showing overlap between transcripts detected with G3BP1 TRIBE-ID and transcripts found in rG4-Seq[4]. **(b)** Venn diagram showing overlap between transcripts detected with G3BP1 TRIBE-ID under PDS treatment and transcripts found in rG4-Seq[4]. **(c)** Scatter plot showing correlation between editing during drug-inhibited PDS and RSVL or EGCG treatments. dS = S(drug) – S(control).

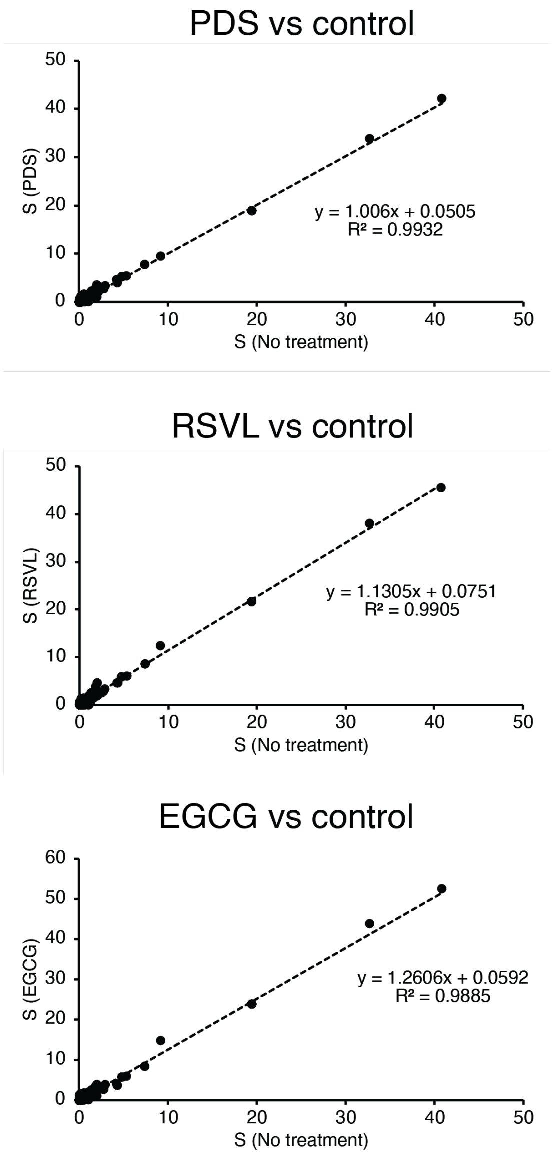


**Supplementary Figure 15**: G3BP1 inhibitors have minimal effect on ADAR activity. Scatter plot showing correlation between background editing (editing present without rapamycin treatment) in drug-treated cells untreated cells.

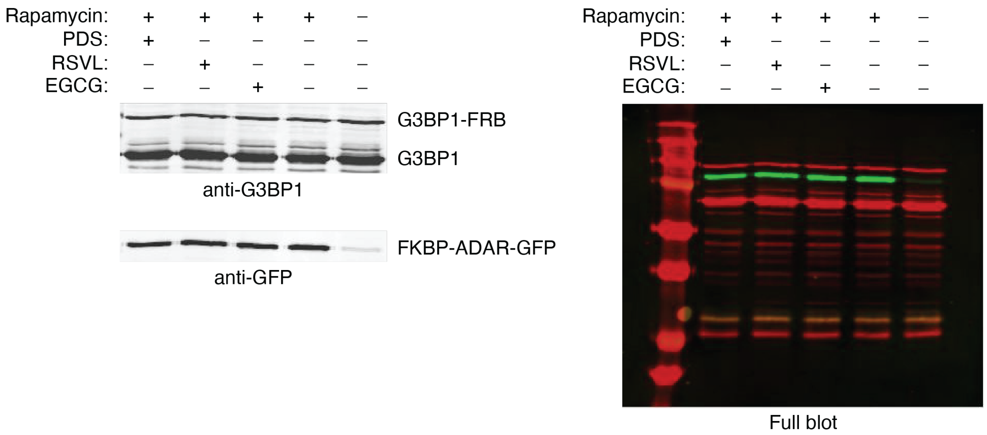


**Supplementary Figure 16**: Co-immunoprecipitation of G3BP1-FRB and FKBP-ADAR in the presence of rapamycin and G3BP1 inhibitors. Full blot shown on the right.

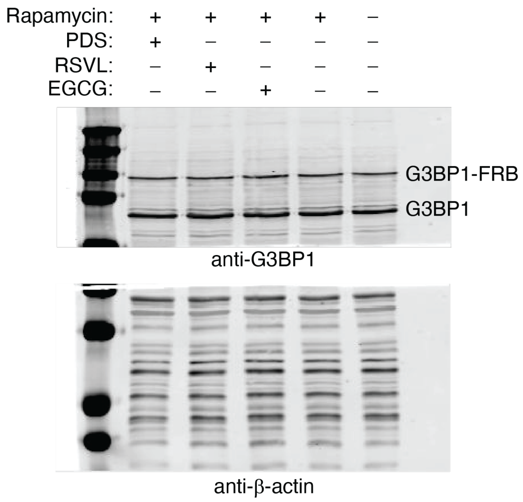


**Supplementary Figure 17**: Western blot analysis of G3BP1 and G3BP1-FRB expression in the presence of G3BP1 inhibitors.

Decreased in PDS

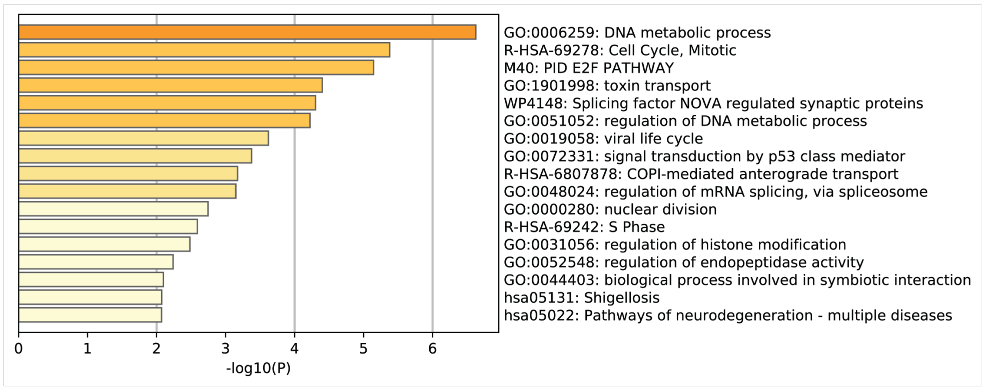


Decreased in RSVL

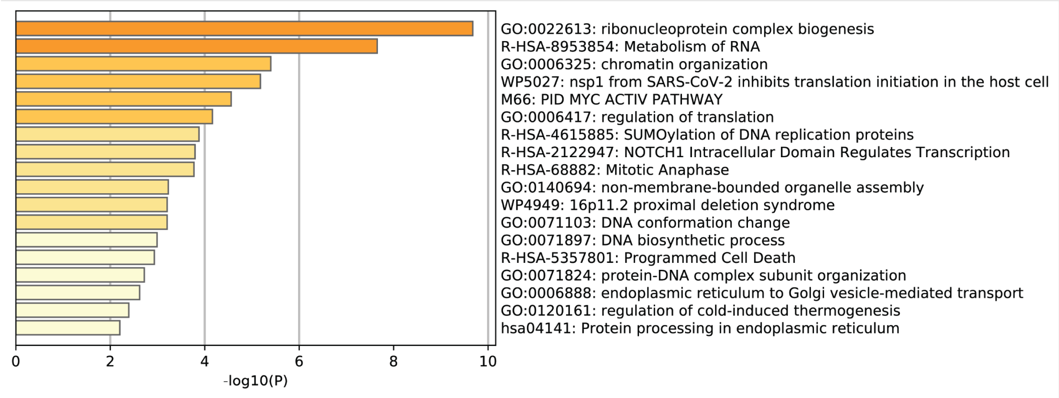


Decreased in EGCG

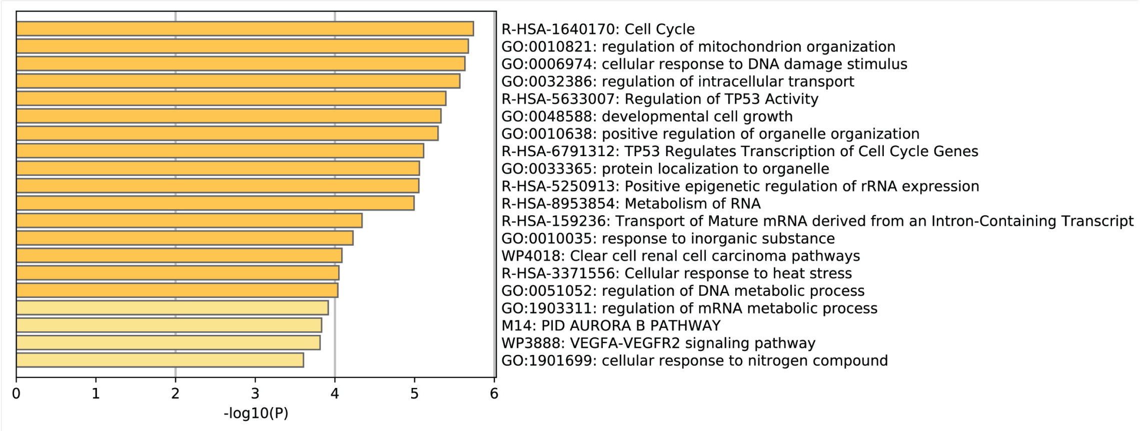


**Supplementary Figure 18**: Top 20 enriched gene ontology terms for G3BP1 TRIBE-ID transcripts with decreased editing in each drug treatment.
